# Assessing the performance of current strain resolution tools on long-read metagenomes

**DOI:** 10.1101/2024.11.20.624313

**Authors:** Ayorinde O. Afolayan, Stefany Ayala Montaño, Ifeoluwa J. Akintayo, Leonardo Duarte dos Santos, Sandra Reuter

**Affiliations:** Institute for Infection Prevention and Control, Medical Centre, University of Freiburg, Freiburg, Germany

**Keywords:** Strain resolution, transmission clusters, long-read sequencing, genomic surveillance, microbial communities, Oxford Nanopore Technology

## Abstract

Recent advances in long-read sequencing-based methods have greatly enhanced genomics and public health applications. However, the challenge of effectively distinguishing strains within microbial communities from clinical samples using these technologies restricts their widespread use. We assessed the strain resolution capabilities of three currently available bioinformatics tools—TRACS, Strainy, and Strainberry—using both mock communities and authentic metagenomic datasets.

Following sample preparation and long-read sequencing using the GridION sequencing platform, raw reads were processed using TRACS, aligning them to a custom reference database, while Strainberry and Strainy mapped reads to metagenome assemblies for strain resolution. Performance on mock microbial community was assessed by comparing predicted microbiota composition to the expected composition, and on both mock and authentic datasets by evaluating strain-resolved genome assemblies. Computational efficiency was measured in terms of task execution time, single-core CPU usage, and physical memory usage.

TRACS demonstrated substantial agreement with the known composition, achieving a median score of 86.7% for *Escherichia coli*-dominant communities and 94.7% for *Klebsiella pneumoniae*-dominant communities. Strainberry and Strainy exhibited improved concordance after excluding strains with a genome size below 1 Mb, thus showcasing comparable performance metrics to TRACS. In mock and real metagenomic datasets, TRACS demonstrated the highest haplotype completeness compared to the other two tools, while Strainy demonstrated the highest haplotype accuracy. All tools were able to allocate strains to their respective transmission clusters (< 20 SNPs), albeit with varying degrees of success. Except for single core CPU usage, TRACS outperformed Strainy and Strainberry in terms of speed and computational efficiency.

Our study underscores the utility of TRACS, Strainy, and Strainberry in resolving strains within microbial communities from clinical samples. TRACS stands out for its better haplotype completeness and computational efficiency, suggesting its potential to streamline advanced genomic analyses and public health initiatives.

## Introduction

Metagenomic sequencing has revolutionised the study of microbial communities, enabling the comprehensive analysis of complex microbial ecosystems within various environments, including the human body [1]. This approach involves the sequencing of genetic material directly obtained from environmental or clinical samples, offering insights into the diverse composition and functional potential of microbial communities [1].

The ability to distinguish between closely related strains within microbial communities from clinical samples holds immense medical significance. Strain-level resolution provides crucial insights into pathogen evolution, transmission dynamics, and virulence variations, particularly essential in infectious disease surveillance and outbreak investigations [2, 3].

Several methods have been employed for strain resolution from metagenomes, primarily relying on short-read sequencing technologies [1, 3, 4]. However, the inherent constraints of short-read sequencing, encompassing incomplete genome coverage, challenges in resolving structural variations and regions with high GC content, and difficulties in handling repetitive sequences, impede accurate strain identification and assembly [3, 5]. Consequently, these constraints present challenges in differentiating closely related strains, curtailing the wider adoption of metagenomic sequencing in clinical and public health settings. In contrast, long-read sequencing technologies, such as those provided by platforms like Oxford Nanopore (ONT) and PacBio, offer extended read lengths and with recently improved error rates (e.g., via R10 chemistry for ONT [6]), they enable more comprehensive genome coverage and better resolution of complex genomic regions [3, 7]. This advancement has the potential to significantly improve the accuracy of strain-level identification from metagenomic data. However, despite their potential, the effective utilization of long-read data for strain resolution necessitates the development of sophisticated bioinformatics tools, which are currently sparse, relatively new, and unproven.

While existing long-read assemblers like Canu [8] and Flye [9] were originally designed for constructing species-level consensus assemblies from long-reads and thus fall short in strain resolution, promising strain resolution tools are emerging. Here, we investigated three currently available tools: Strainberry, TRACS, and Strainy. Strainberry is a bioinformatics pipeline adept at strain resolution in low-complexity metagenomes [10]. It employs variant calling from long-read sequence data, haplotype phasing, and grouping reads into clusters corresponding to strains, followed by strain assembly. TRACS [11] infers pairwise transmission from single genomes or metagenomic data using a kmer-based approach. It calculates SNP distances across all strains by aligning sequencing reads to a custom or Genome Taxonomy Database (GTDB)/sourmash database [12], grouping strains into putative transmission clusters based on pairwise SNP and transmission distance estimates. The Strainy pipeline [13] aligns reads against metagenome-assembled genome (MAG) contigs. It identifies regions with collapsed strains and phases them into strain-resolved assemblies, referred to as haplotigs, for the corresponding MAG species. While these tools strive to harness long-read data for strain resolution in metagenomic samples, a critical assessment of their performance, particularly in complex microbial communities, is indispensable. Additionally, a comparative evaluation of their computational efficiency, encompassing factors like execution time and resource utilization, is pivotal for determining their practical viability in diverse settings.

In this study, we aim to evaluate the strain resolution capabilities of the aforementioned tools in defined mock microbial communities as well as real microbial communities. By assessing their strain identification accuracy, efficiency, and robustness in the context of different bacterial strains and community compositions, we seek to address the limitations associated with strain resolution from metagenomes and further advance the application of long-read sequencing in clinical and public health contexts.

## Methodology

### Ethical Approval

As the study did not involve the use of associated epidemiological and clinical data, no ethical approval was required.

### Construction of Defined Mock Microbial Communities

Pure cultures of reference *Escherichia coli* strains (n = 10) and *Klebsiella pneumoniae* strains (n = 10) were cultured on blood agar at 37 °C for 18 – 24 hours. Reference strains used in this study were initially isolated during routine rectal and nasal screening of neonates within the neonatal intensive care unit (NICU) at the University of Freiburg Medical Centre, Freiburg, Germany. Previously, genomic DNA was extracted from these strains using the High Pure PCR Template Preparation Kit (Roche) according to manufacturer’s protocol. DNA libraries were then prepared with the Nextera DNA Flex Library Preparation Kit (Illumina). Paired-end sequencing (2×150 reads, 300 cycle v2) was conducted on an Illumina MiSeq platform (Illumina). Following the re-extraction of DNA from each reference strain, batches of mock microbial communities were defined by combining known proportions of the reference strains’ DNA (Table 1; refer to the “DNA Extraction, library preparation, and Metagenomic Sequencing” step for more details).

**Table 1:** Summary of taxonomy identity, sequence types and proportion of strains making up the mock microbial community.

| Metagenome ID | Species | Sequence Type | Proportion |
| --- | --- | --- | --- |
| 13723_17053_12926 | *K. pneumoniae* | 11, 147, 258 | 10:25:65 |
| 15240_16126_5925 | *K. pneumoniae* | 258, 11, 147 | 20:30:50 |
| 16464_Kpn01856 | *K. pneumoniae* | 307 | 30:70 |
| 16902_13733 | *K. pneumoniae* | 258, 307 | 30:70 |
| 17266_Kpn01933_Kpn01518 | *K. pneumoniae* | 258 | 20:30:50 |
| 760592820_Kpn01890_Kpn01915 | *K. pneumoniae* | 307 | 10:25:65 |
| Kpn01412_Kpn01528 | *K. pneumoniae* | 258 | 40:60 |
| Kpn01412_Kpn01886 | *K. pneumoniae* | 258, 307 | 50:50 |
| Kpn01528_Kpn01886 | *K. pneumoniae* | 258, 307 | 40:60 |
| Kpn01886_13733 | *K. pneumoniae* | 307 | 30:70 |
| Eco04708_Eco04812 | *E. coli* | 73 | 40:60 |
| Eco04711_Eco04737_Eco04896 | *E. coli* | 141, 73, 10 | 20:30:50 |
| Eco04714_Eco04708 | *E. coli* | 141, 73 | 50:50 |
| Eco04714_Eco04749 | *E. coli* | 141 | 50:50 |
| Eco04720_Eco04711_Eco04831 | *E. coli* | 141 | 20:30:50 |
| Eco04720_Eco04799 | *E. coli* | 141, 73 | 30:70 |
| Eco04737_Eco04799 | *E. coli* | 73 | 30:70 |
| Eco04749_Eco04812 | *E. coli* | 141, 73 | 40:60 |
| Eco04800_Eco04851_Eco04723 | *E. coli* | 141 | 10:25:65 |
| Eco04831_Eco04883_Eco04834 | *E. coli* | 141, 1193, 131 | 10:25:65 |

### Collection of Samples from Neonates at the Neonatal Intensive Care Unit

Rectal and nasal samples were routinely collected from neonates undergoing admission to the NICU (Neonatal Intensive Care Unit) within the framework of a study tracking the acquisition of pathogens in real time (TAPIR). This project seeks to investigate the emergence, spread, and dynamics of healthcare-associated, high-risk bacterial clones within both local and regional contexts. From this pool of samples, we selected a total of 36 samples to evaluate the performance of strain resolution tools where *K. pneumoniae* and/or *E. coli* were identified as the most prevalent or second most prevalent strains during regular metagenomic analysis or were identified during bacterial culture. Additionally, we included samples that tested negative for microbiological growth, as well as samples suspected to contain multiple strains of the species (*E. coli* or *K. pneumoniae*) within the metagenome.

### DNA Extraction, Library Preparation, and Sequencing

Samples derived from the collection of rectal and nasal swabs were enriched with BHI overnight at 37°C, 210 rpm in a proportion 1:2 (sample:media). Following enrichment, four types of solid media were incubated (MacConkey, Blood Agar, CNA, and Chromogenic Agar) at 37°C for 24 hours. For each distinct colony morphology observed on MacConkey, a pure culture was isolated. Microbial species were presumptively identified from these pure cultures using MALDI-TOF MS. DNA was extracted using the High Pure PCR Template Preparation Kit (Roche) following the manufacturer’s instructions. DNA libraries were prepared using the Nextera DNA Flex Library Preparation Kit (Illumina), and paired-end sequencing (2×150 bp reads, 300 cycles) was performed on an Illumina MiSeq platform. The rest of the BHI enrichment was processed for the removal of human DNA by a lysis protocol [14] that involved the re-suspension of the pellet with 1 mL of clean dH_2_O at RT for 5 minutes and a final resuspension in 200 μL PBS 1X, 5 uL lysozyme, and 5 uL lysostaphin. Microbial DNA extraction from the 36 samples and from pure cultures of reference strains derived from these samples were performed using the High Pure PCR Template Preparation Kit (Roche Molecular Diagnostics, Mannheim, Germany), following the manufacturer’s protocol. Following DNA extraction, synthetic genomic pools of defined mock microbial communities were prepared by combining up to three Gram-negative bacteria of the same species but with varying sequence types—first using different sequence types and then using the same sequence type—in varying proportions (Table 1). Libraries for GridION sequencing were prepared using a ligation sequencing kit (SQL-LSK109; ONT, Oxford, UK) and native barcoding kits (EXP-NBD104 & EXP-NBD114) for R9 Chemistry, as well as the SQK-NBD114-96 barcoding kit for R10 chemistry, following the manufacturer’s protocol. The following optional steps were also carried out:

i. DNA input and other measurements were calculated based on the assumption that 1 fmol is equivalent to 5 ng of DNA. Thus, for the final R10 library, 100 ng corresponded to 20 fmol.
ii. In the R10 protocol,

a. the initial volume per sample was 11 µL without water dilution (for clinical samples).
b. optional NEBNext FFPE DNA Repair and DNA Control Sample were omitted during DNA repair and end-prep.
c. Bovine Serum Albumin (BSA) was used to improve the sequencing performance as recommended in the protocol.

Final libraries were quantified using Qubit 4 Fluorometer, loaded onto a FLO-MIN106 R9.4 SpotON flow cell (for defined mock communities) and a FLO-MIN114 R10.4 flow cell (for defined mock communities and real-world microbial communities), and sequenced on a GridION X5 Mk1 sequencing platform. Sequence data acquisition, real-time base-calling, and demultiplexing of barcodes were conducted using the graphical user interface MinKNOW (v23.11.7) and the dorado basecall server (v7.2.13).

### Read Pre-processing and Removal of Human DNA contaminants

Sequenced data underwent pre-processing using Porechop (v0.2.4; https://github.com/rrwick/Porechop) to trim off adapters. Trimmed reads were filtered using Filtlong (v0.2.1; https://github.com/rrwick/Filtlong) with a minimum read length threshold (“`--min_length`”) of 1000. Afterwards, reads were mapped to the human genome (GRCh38) using Minimap2 (v2.24; using the “`map-ont`” parameter) [15] and human DNA contaminants were removed from reads using Samtools (v1.14) [16]. These steps, including read trimming, read filtration, and human DNA removal, were executed using custom Nextflow pipelines [17] (https://github.com/ayoraind/ONT_adapter_removal_and_read_filtration; https://github.com/ayoraind/hDNA_removal_and_mapping_stats; see “code availability” section of the methodology).

### Microbial Species Abundance Estimation and Visualization of Real-world Metagenomic Datasets

Kraken reports were generated from the analysis of metagenomic sequence data, using Kraken (v1.1.1) [18] and the minikraken 4 GB reference database, implemented within a custom Nextflow pipeline (https://github.com/ayoraind/kraken). Microbial species abundance was estimated from kraken output using Bracken (v2.8) [19], which was incorporated into a custom Nextflow pipeline (https://github.com/ayoraind/bracken). Microbial species abundance profiles were visualized in R, using Tidyverse (v2.0.0) [20], scales [21], and ggthemes [22].

### TRACS

To evaluate the performance of TRACS [12] on mock datasets, alignments were generated using the `“tracs align”` command (https://gtonkinhill.github.io/tracs/#/alignment; https://github.com/ayoraind/nxf_Tracs) on each metagenomic sequence data. A custom database of references, given the known and defined components of the mock community (Table 1), was employed. Matches to the reference were identified from the suffix of the output fasta file (e.g., the “T183-28-A-Kpn-231026_contigs_filtered” in T100-12-A_G230331_posterior_counts_ref_T183-28-A-Kpn-231026_contigs_filtered.fa).

Completeness and accuracy statistics were computed using metaQUAST [23], incorporating the “--unique-mapping” and “--reuse-combined-alignments” options to ensure that a single contig only contributed to the completeness of one reference strain.

For real-world metagenomic datasets, TRACS’s performance was tested by generating alignments with the `“tracs align”` command on each metagenomic sequence data. A custom database of *K. pneumoniae* and *E. coli* references was utilized (see sub-section “DNA Extraction, Library Preparation, and Sequencing” for details on the source of reference genomes), selected from a representative of each cluster in our collection using ska (v1.0) [24]. Like the mock analysis, the quality of the assemblies was assessed using metaQUAST.

### Strainberry

Sequence data were assembled using metaFlye (Flye (v2.9.1) with the “`--meta`” parameter). These reads were mapped to the associated assemblies using Minimap2 to generate long-read mapping files. Strain-resolved assemblies were generated from the strain-oblivious assembly (Flye) and long-read mapping files using Strainberry (v1.1). Assembly quality and statistics were computed using metaQUAST (with `“--unique-mapping`“ and `“--reuse-combined-alignments”` options) and the Strainberry assembly statistics python script (https://github.com/rvicedomini/strainberry-analyses/blob/main/workflow/scripts/assembly_stats.py). The reference database for testing the strain resolution performance of Strainberry on mock communities and real microbial communities contains the same references used to test the strain resolution performance of TRACS.

### Strainy

Sequence data were assembled using metaFlye (Flye v2.9.1) with the metaFlye polishing procedure disabled, as recommended. Strainy (pre-release version) takes the assembly graph files and the sequence data as input to generate strain-resolved assemblies. Assembly quality was assessed using the same tools and reference databases used for assessing the quality of Strainberry assemblies.

### Tree Construction, Concordance Analysis, and Visualization

A core genome phylogenetic tree was constructed from core genes previously generated from pangenome analysis [25, 26]. The level of concordance between the known composition of the mock community (ground truth) and the observed composition following strain resolution using the three tools, along with additional metrics like F1 score, sensitivity, and specificity, was assessed using the epiR package (v2.0.60) [27]. The interactive tree of life (iTOL) annotation files were generated using the itol.toolkit (v1.1.5) [28] in R (v4.2.2) [29]. The phylogenetic tree and data showcasing the performance of the three strain resolution tools were visualized in iTOL [30]. Plots were visualized using ggplot from Tidyverse (v2.0.0) [20]. Upset plots [31] were visualized using UpSetR (v1.4.0), ggupset (v0.3.0), and ComplexUpset (v1.3.5).

### Computational Performance

The computational efficiency of the strain resolution tools was evaluated by analyzing the pipeline information generated through their respective Nextflow pipelines. The pipelines were run on an Ubuntu Linux distribution (v.20.04) with ×86_64 architecture.

### Code Availability

The codes for the specified processes were implemented in Nextflow pipelines [17] and are accessible at the following links:

i. Read trimming and filtering: https://github.com/ayoraind/ONT_adapter_removal_and_read_filtration
ii. Human DNA contaminant removal: https://github.com/ayoraind/hDNA_removal_and_mapping_stats
iii. Kraken: https://github.com/ayoraind/kraken
iv. Bracken: https://github.com/ayoraind/bracken
v. TRACS: https://github.com/ayoraind/nxf_Tracs
vi. Flye: https://github.com/ayoraind/flye
vii. Strainberry: https://github.com/ayoraind/strainberry
viii. Strainberry accessories: https://github.com/ayoraind/strainberry-accessories
ix. Strainy: https://github.com/ayoraind/strainyMAG
x. Core genome phylogeny: https://github.com/ayoraind/genome-annotation-and-pangenome-analysis

## Results

### Strain resolution tools show comparable performance on mock microbial communities

We assessed the performance of TRACS, Strainberry, and Strainy by comparing the composition of the known mock communities to the observed composition following strain resolution. We found no significant difference in the performance of the three strain resolution tools using standard metrics, particularly when contigs smaller than 1 Mb in genome size were excluded from the resolved mock community using Strainberry and Strainy (Figure 1A and 1B). Moreover, our analysis revealed that the performance of these tools was consistent across different flow cell chemistries and dominant species within the mock community (Figure 1A and 1B).

**Figure 1:**
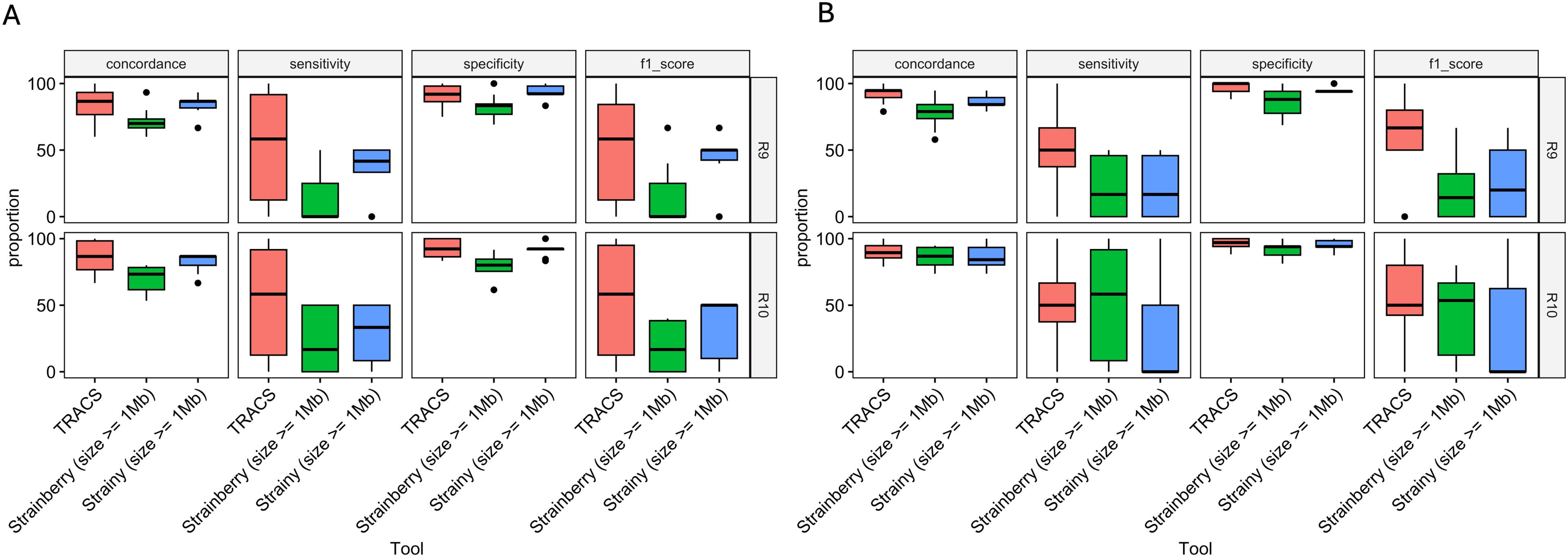
Comparative analysis of strain resolution tool performance on *E. coli*-dominant (A) and *K. pneumoniae*-dominant (B) mock communities, stratified by the flow cell chemistry employed. Evaluation metrics include concordance, specificity, sensitivity, and F1 score, benchmarked against the known composition of the mock microbial community.

### Strain resolution performance is dependent on the SNP distance between strains within the mock community

Our investigation into strain resolution dynamics revealed a consistent pattern: when the expected strains within a mock community share the same sequence type but are several SNPs apart, or if they belong to different sequence types, strain resolution tools typically identify the exact or the closest matching strain from the reference database in most cases (Figure 2, Figure S1). This holds true across various flow cell chemistries (Figure 2). However, we observed that Strainberry tended to overestimate the richness and diversity of strains within the mock community, while Strainy tended to underestimate this, especially when the strains belonged to the same sequence type (Figure 2, Figure S1). Furthermore, Strainberry and Strainy sometimes identify strains that are not the closest match to the expected strain. For instance, in mock community comprising two *E. coli* strains (ratio 2:3) of the same sequence type (figure 2 section A), Strainberry overestimated the strain count within the mock metagenome, using the R10 chemistry (see “Observed_M1_SB_1Mb_R10”). In this case, both the expected strains and strains from the same cluster were identified, rather than just the expected strains or those within the same cluster. In a mock community consisting of two strains (ratio 1:1) from different sequence types (Figure 2, Section B), Strainberry once again overestimated the number of strains detected, irrespective of the flow cell chemistry used. Conversely, while Strainy identified two strains, one of them was incorrectly classified, as it belonged to a different cluster (albeit the same sequence type) compared to the expected strain. When the expected number and diversity of expected strains within the mock microbial community increased to three (ratio 2:3:5; Figure 2 section C), even TRACS surprisingly overestimated the number of strains within the mock microbial community, using the R9 chemistry. However, the overestimation issue was resolved with the R10 chemistry. In this scenario, Strainberry performed accurately by identifying the correct strains or those within the same cluster. Strainy, on the other hand, underestimated the number of strains present, regardless of the flow cell chemistry used. When the mock community comprise up to three strains belonging to the same sequence type (ratio 2:3:5; Figure 2 section D), TRACS was able to resolve two of these, regardless of the flow cell chemistry used. Although Strainberry resolved three strains, one of the strains was wrongly identified, as it did not belong to the same cluster as the expected strain. Strainy underestimated the number of strains within the mock community, as it was only able to resolve one of the three expected strains alone.

**Figure 2:**
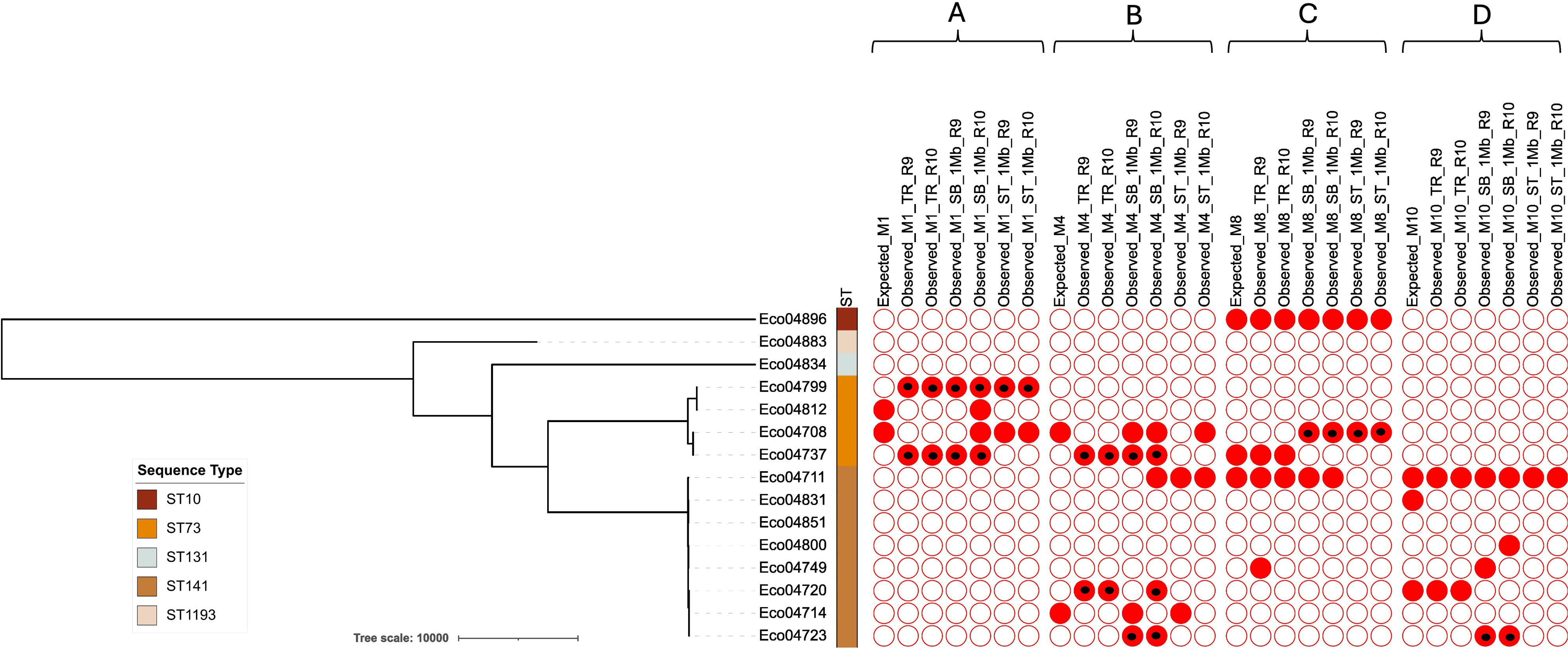
Maximum-likelihood phylogenetic tree of core genomes depicting microbial strains within the mock microbial community. Known *E. coli* strain identities are juxtaposed with strains detected by the three strain resolution tools. Annotations on the right side of the tree denote strains’ sequence type as well as expected and predicted mock composition, with red-shaded circles indicating presence and empty circles indicating absence. Black dots within red-shaded circles signify strains belonging to the same cluster (SNP < 20) as the expected strain identity. Section A and section B refer to mock community comprising up to two strains belonging to the same sequence type and different sequence types, respectively. Section C and section D refer to mock community comprising up to three strains belonging to a different sequence type and same sequence types, respectively.

In a mock community comprising two *K. pneumoniae* strains (ratio 1:1) from different sequence types (Figure S1, Section A), TRACS correctly identified the cluster for one of the strains. However, for the second strain, TRACS failed to identify both the strain and its correct cluster, even though the strain identified belonged to the same sequence type as the expected strain. Strainberry accurately identified both expected strains but overestimated the strain count. On the other hand, Strainy underestimated strain diversity, detecting only one of the two possible strains, and misidentified it as belonging to a different cluster, despite it being of the same sequence type as the expected strain. In a mock community consisting of two strains (ratio 2:3) from the same sequence type (Figure S1, Section B), both TRACS and Strainberry detected only one of the strains. Strainy identified a strain that shared the same sequence type as one of the expected strains but did not belong to the same cluster. In a mock community consisting of three strains (ratio 2:3:5) from different sequence types (Figure S1, Section C), TRACS underestimated the number of strains, although it correctly detected those identified. Once again, Strainberry overestimated the strain count within the community, irrespective of the flow cell chemistry used. Strainy underestimated the strain count and was able to predict only the sequence type of the identified strains, without accurately identifying the expected strain or cluster. Finally, when the mock community comprise up to three strains (ratio 10:25:65) belonging to the same sequence type (Figure S1 section D), all three tools resolved only one of the three strains.

### TRACS demonstrates higher haplotype completeness, while Strainy exhibits superior haplotype accuracy in mock microbial community datasets

We employed metaQUAST to assess the completeness and accuracy of strain assemblies derived from the mock metagenome dataset (Figure S2 and Figure S3). Across all genomes within the metagenome, the genome fraction—indicating the proportion of reference genomes assembled by each strain resolution tool—varied notably: TRACS achieved a range of 0% to 95.8%, Strainberry ranged from 0% to 97.7%, and Strainy exhibited a range of 1.31% to 84.7% when utilizing R9 flow cell chemistry on *E. coli*-dominant mock communities. Following the transition to R10 flow cell chemistry, we observed a slight improvement in genomic fraction for TRACS (0% – 99.6%) and Strainberry (0.895% – 98%), albeit a reduction for Strainy (0.582% – 81.9%). It is worth noting that the absence of the expected reference genome in the metagenome always yielded a genome fraction of 0%. In mock communities comprising up to two *E. coli* strains belonging to either the same sequence type or different sequence types, the highest genome fraction was observed in TRACS-resolved metagenomes, using the R9 and R10 flow cell chemistry (Figure 3A). A similar trend was noted in mock community comprising up to three *E. coli* strains belonging to the same sequence type. However, in mock community comprising up to three strains belonging to different sequence types, the genome fractions were comparable between TRACS-resolved metagenomes and Strainberry-resolved metagenomes. In mock communities comprising up to two or three *K. pneumoniae* strains belonging to the same sequence types, the highest genome fraction was observed for TRACS-resolved metagenomes, using R9 flow cell chemistry (Figure 3B). Nevertheless, in mock communities with two or three strains, whether from the same or different sequence types, genome fractions generally decreased across all tools when using R10 flow cell chemistry, with Strainberry showing a slight advantage, yielding the highest genome fraction among the three tools tested (Figure 3B).

**Figure 3:**
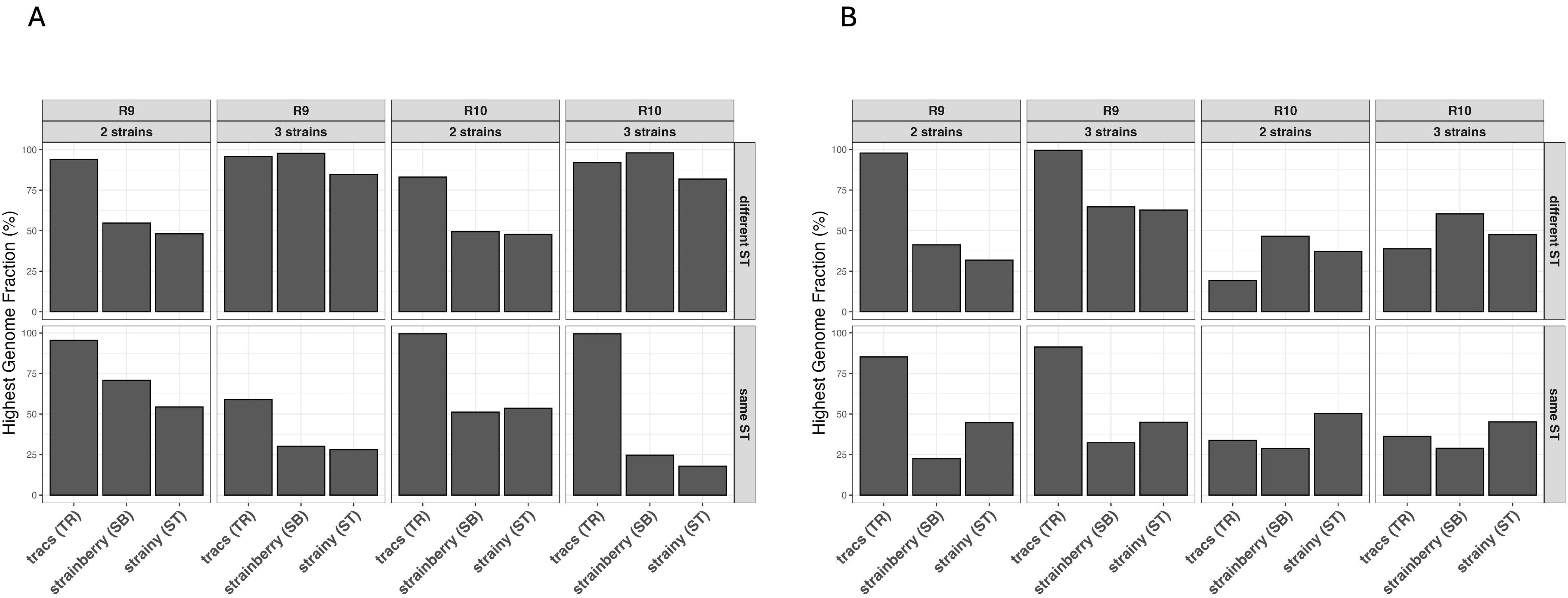
Highest assembled strain genome fraction in *E. coli* and *K. pneumoniae*-dominant mock metagenome datasets.

Regarding misassemblies, Strainy showed the lowest occurrence, with a maximum of 122 misassemblies detected, while Strainberry exhibited the highest, with up to 256 misassemblies observed using R9 chemistry (Figure S1). Predominantly, misassemblies were noted in mock communities containing up to three strains of diverse sequence types. Furthermore, Strainy consistently displayed the least number of misassemblies, with only 2 identified, in mock communities comprising up to three strains of the same sequence type, contrasting with Strainberry (4) and TRACS (21). This trend persisted with R10 chemistry, although with varying numbers, except for Strainy and Strainberry exhibiting an equal lowest count of misassemblies (n = 4).

When compared to *E. coli*-dominant mock communities, TRACS exhibited the highest number of misassemblies (n = 76) in communities containing up to two *K. pneumoniae* strains of different sequence types. Conversely, Strainberry (n = 28) and Strainy (n = 26) showed fewer misassemblies, particularly in communities comprising up to three strains of the same sequence type. Notably, Strainy demonstrated the lowest misassembly count (n = 12) in communities containing up to two strains, even with R10 chemistry, indicating a superior performance over TRACS and Strainberry.

### Strain-level resolution details of the three tools on Real-world Metagenomic datasets

Bracken analysis revealed the presence of *K. pneumoniae* and *E. coli* across all 36 real-world metagenomes. Among these, TRACS, Strainy, and Strainberry resolved *E. coli* strains and/or *K. pneumoniae* strains in 34, 27, and 28 metagenomes, respectively. Additionally, analysis of short-read sequence data from pure culture resulted in the identification of *E. coli* and/or *K. pneumoniae* strains in 32 of the 36 samples. Figure 4 illustrates the distribution of *K. pneumoniae* and *E. coli* species within the 36 authentic communities assigned for benchmarking, with species constituting less than 1% abundance categorized as “Other”.

**Figure 4:**
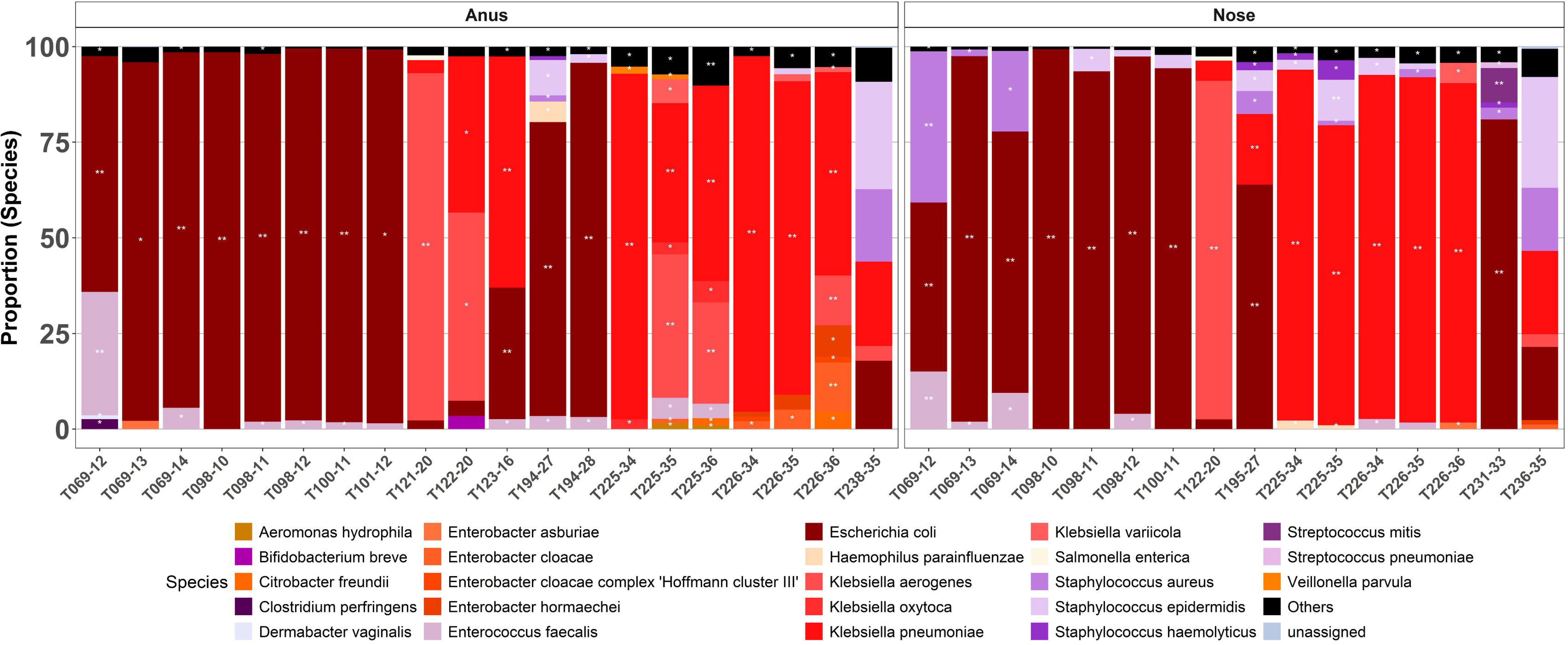
Taxonomic profile of real-world datasets stratified by sample collection site. Double white stars on stacked bars represent species with ≥10,000 reads assigned to them, while a single white star indicates species with 1,000 to 10,000 reads. Species with less than 1,000 reads are denoted by no star.

In both microbiologically negative samples, all strain resolution tools correctly reported the absence of *E. coli* and *K. pneumoniae*. Bracken, however, assigned a very low number of reads to these species. Furthermore, TRACS, Strainy, Strainberry, and Illumina data analysis agreed on the presence and identity of species present within one other metagenome (Table 2). In general, whenever *E. coli* and/or *K. pneumoniae* strains were identified through short-read analysis, TRACS detected the same strain (or at least one of the two strains identified by Illumina) in all but one metagenome. For this particular metagenome, further investigation revealed that the TRACS custom database did not contain a reference genome corresponding to the sequence type or cluster of the strain identified by Illumina. Conversely, Strainberry and Strainy either detected a different strain or failed to detect any strain in 9 and 10 metagenomes, respectively, where short-read sequence data analysis confirmed the presence of at least one *E. coli* or *K. pneumoniae* strain. Interestingly, in four of the 36 metagenomes (T069-14-B, T098-12-B, T101-12-A, & T123-16-A), at least one of the three tools (usually TRACS) were able to detect the presence of the dominant *E. coli* strain in cases where short-read sequencing of pure cultures did not yield any results (Table 4).

**Table 2:** Similarities between resolution and taxonomic profile calls of TRACS, Strainy, and Strainberry. The last two samples (indicated with the * sign) are microbiologically negative samples. ND is an acronym for “Not detected”.

| Sample | Species<br>(Bracken) | Reads<br>(n) | Reads<br>(%) | Strain id<br>(Illumina) | Strain id<br>(TRACS) | Strain id<br>(Strainy) | Strain id<br>(Strainberry) |
| --- | --- | --- | --- | --- | --- | --- | --- |
| T231-33-B | *E. coli* | 111344 | 80.97 | EcST14 | EcST14 | EcST14 | EcST14 |
|  | *K. pneumoniae* | 227 | 0.17 | ND | ND | ND | ND |
| T236-35-B* | *E. coli* | 282 | 19.10 | ND | ND | ND | ND |
|  | *K. pneumoniae* | 322 | 1.82 | ND | ND | ND | ND |
| T238-35-A* | *E. coli* | 397 | 17.88 | ND | ND | ND | ND |
|  | *K. pneumoniae* | 489 | 22.03 | ND | ND | ND | ND |

We observed several differences in strain resolution by the three tested tools. For instance, in three metagenomes where TRACS resolved either one or both of *E. coli* and *K. pneumoniae* strains, Strainy and Strainberry were unable to resolve strains of either species (Table 3). In two of the three metagenomes containing *E. coli* or *K. pneumoniae* strains, analysis of short-read sequence datasets was consistent with TRACS’s results. However, in the third metagenome (T122-20-N) where TRACS resolved both *E. coli* and *K. pneumoniae* strains, the short-read sequence analysis initially provided no insights. Upon re-culturing the sample, a *K. pneumoniae* strain, belonging to the same sequence type identified by TRACS, was confirmed (Table 3). Notably, there was no instance where Strainberry and/or Strainy resolved strains from metagenomes that TRACS failed to resolve (Table 4).

**Table 3:**
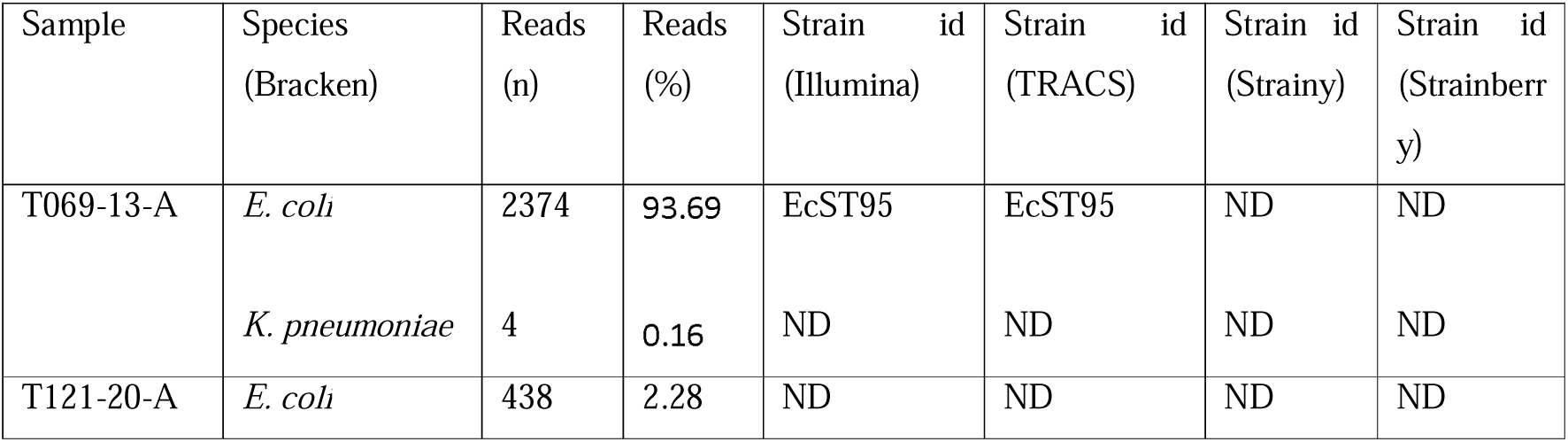
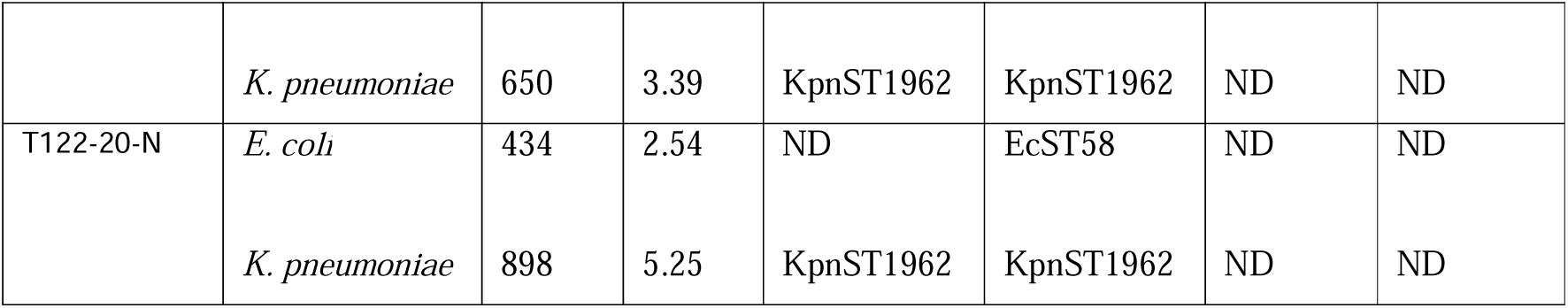
Information on Metagenomes resolved by TRACS, but not resolved by Strainberry and Strainy. ND is an acronym for “Not detected”.

| Sample | Species<br>(Bracken) | Reads<br>(n) | Reads<br>(%) | Strain id<br>(Illumina) | Strain id<br>(TRACS) | Strain id<br>(Strainy) | Strain id<br>(Strainberry) |
| --- | --- | --- | --- | --- | --- | --- | --- |
| T069-13-A | *E. coli* | 2374 | 93.69 | EcST95 | EcST95 | ND | ND |
|  | *K. pneumoniae* | 4 | 0.16 | ND | ND | ND | ND |
| T121-20-A | *E. coli* | 438 | 2.28 | ND | ND | ND | ND |
|  | *K. pneumoniae* | 650 | 3.39 | KpnST1962 | KpnST1962 | ND | ND |
| T122-20-N | *E. coli* | 434 | 2.54 | ND | EcST58 | ND | ND |
|  | *K. pneumoniae* | 898 | 5.25 | KpnST1962 | KpnST1962 | ND | ND |

In six examined metagenomes, TRACS uniquely identified the co-existence of two distinct *K. pneumoniae* strains (KpnST250, KpnST1471-1LV; 20437 – 97929 reads) alongside one *E. coli* strain (EcST501; 235 – 1320 reads) (Figure 4a). This specific combination of strains was not detected by any other tool (Figure 5a; Table 4). Instead, Strainberry detected a single *K. pneumoniae* strain (KpnST1471-1LV) in five of the six metagenomes, with two *K. pneumoniae* strains (KpnST250, ST1471-1LV) found in one metagenome. Strainy detected a single *K. pneumoniae* strain (KpnST1471-1LV) in four out of the six metagenomes, with two *K. pneumoniae* strains (KpnST250, KpnST1471-1LV) identified in one metagenome. Interestingly, in one of the six metagenomes, despite a substantial number of reads assigned to *K. pneumoniae* (20437 reads), Strainy failed to detect either *K. pneumoniae* or *E. coli* strains. Short-read Illumina sequencing of pure cultures isolated from clinical samples confirmed the presence of a KpnST250 strain in two metagenomes and a KpnST1471-1LV strain in four metagenomes (Table 4).

**Figure 5:**
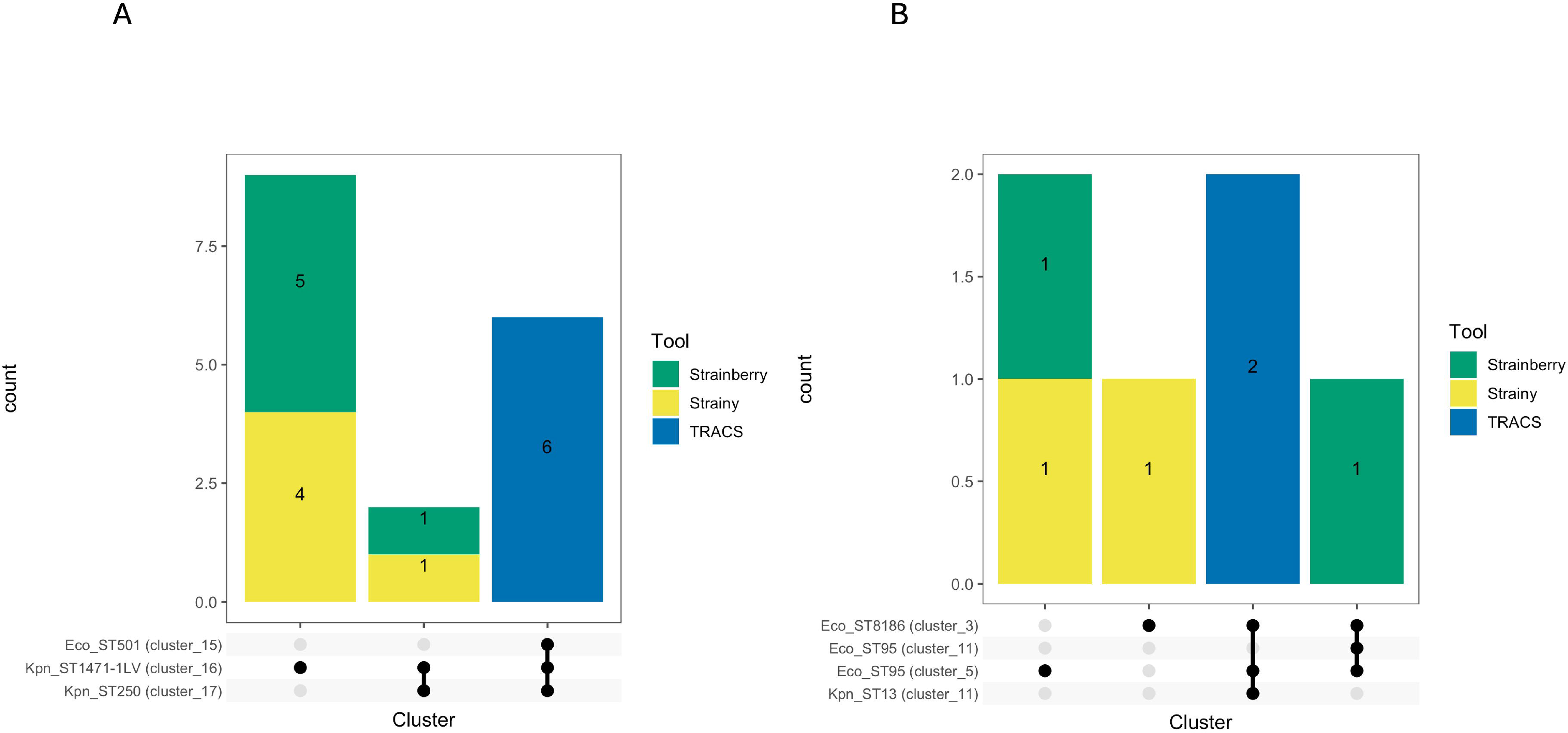
Upset plot illustrating combinations of strains identified/resolved by TRACS (blue colour), Strainy (yellow colour), and Strainberry (green colour) within real-world metagenomes, where TRACS identified one *E. coli* strain and two *K. pneumoniae* strains (A), or where TRACS identified two E. coli strains and one K. pneumoniae strain (B). The main bar chart displays the frequency of metagenomes per strain combination, arranged in descending order. Side bars indicate the number of metagenomes hosting each named strain. Dots and connecting lines at the base of the main bar chart represent the combination of *E. coli* and/or *K. pneumoniae* strains.

**Table 4:**
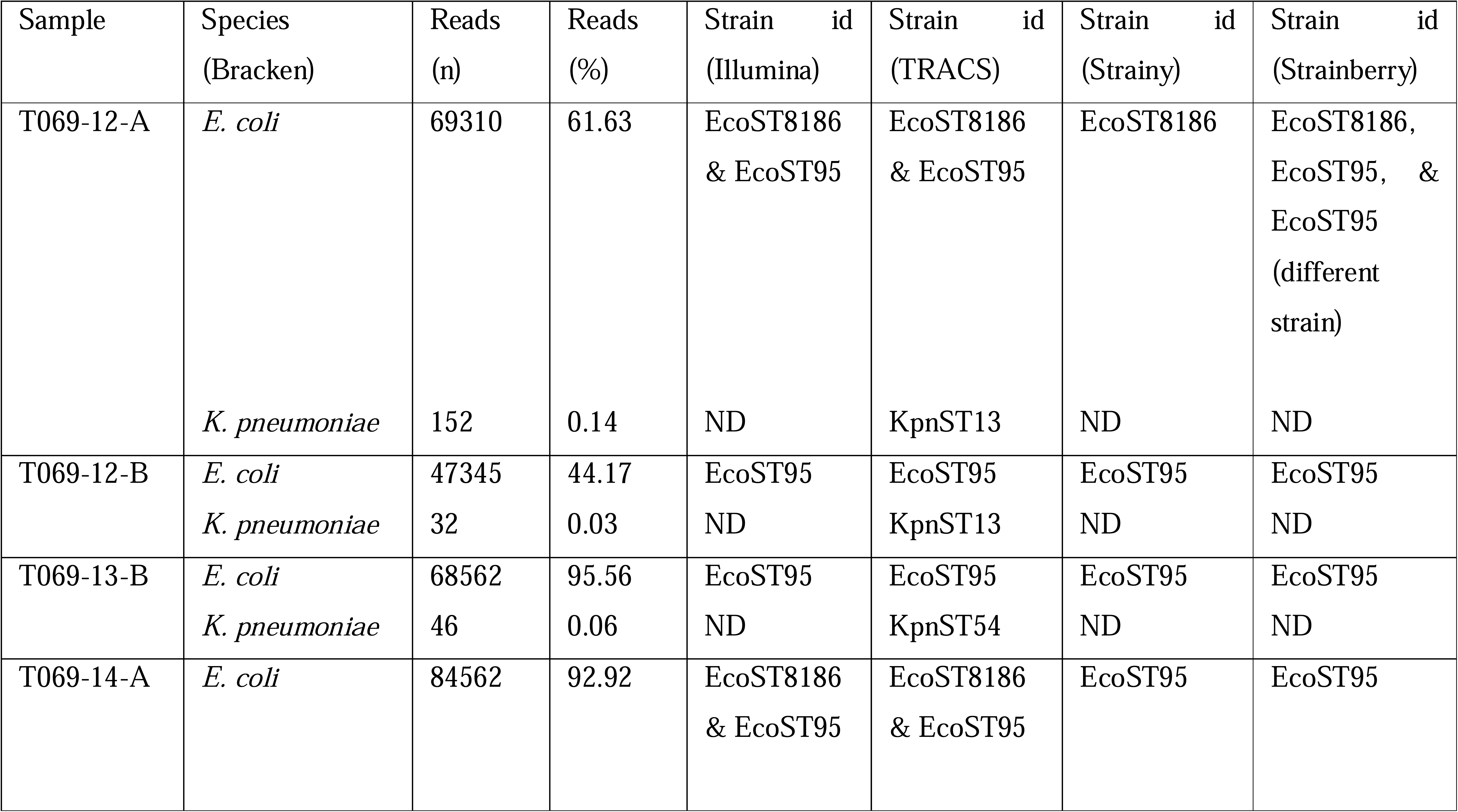
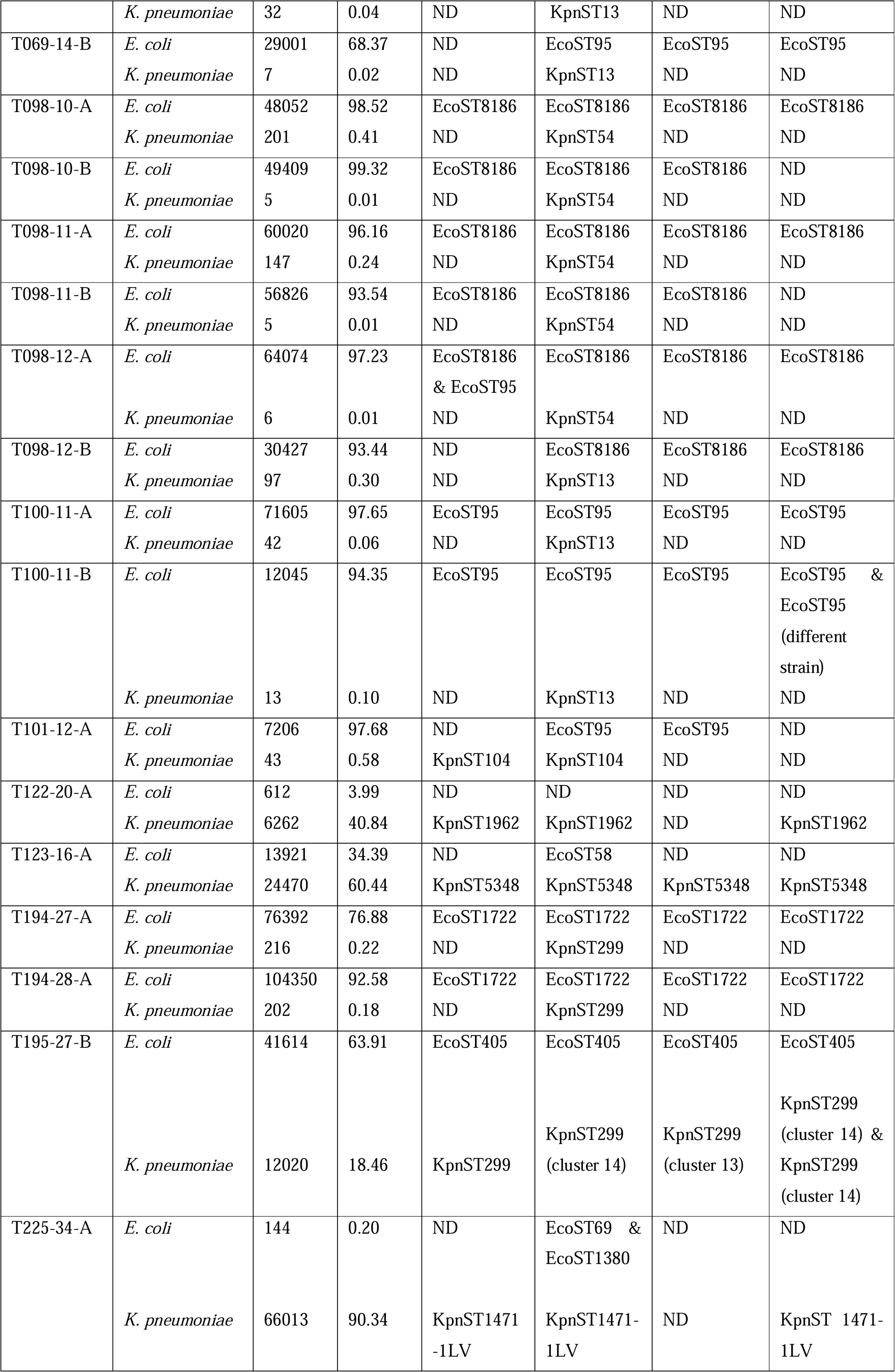
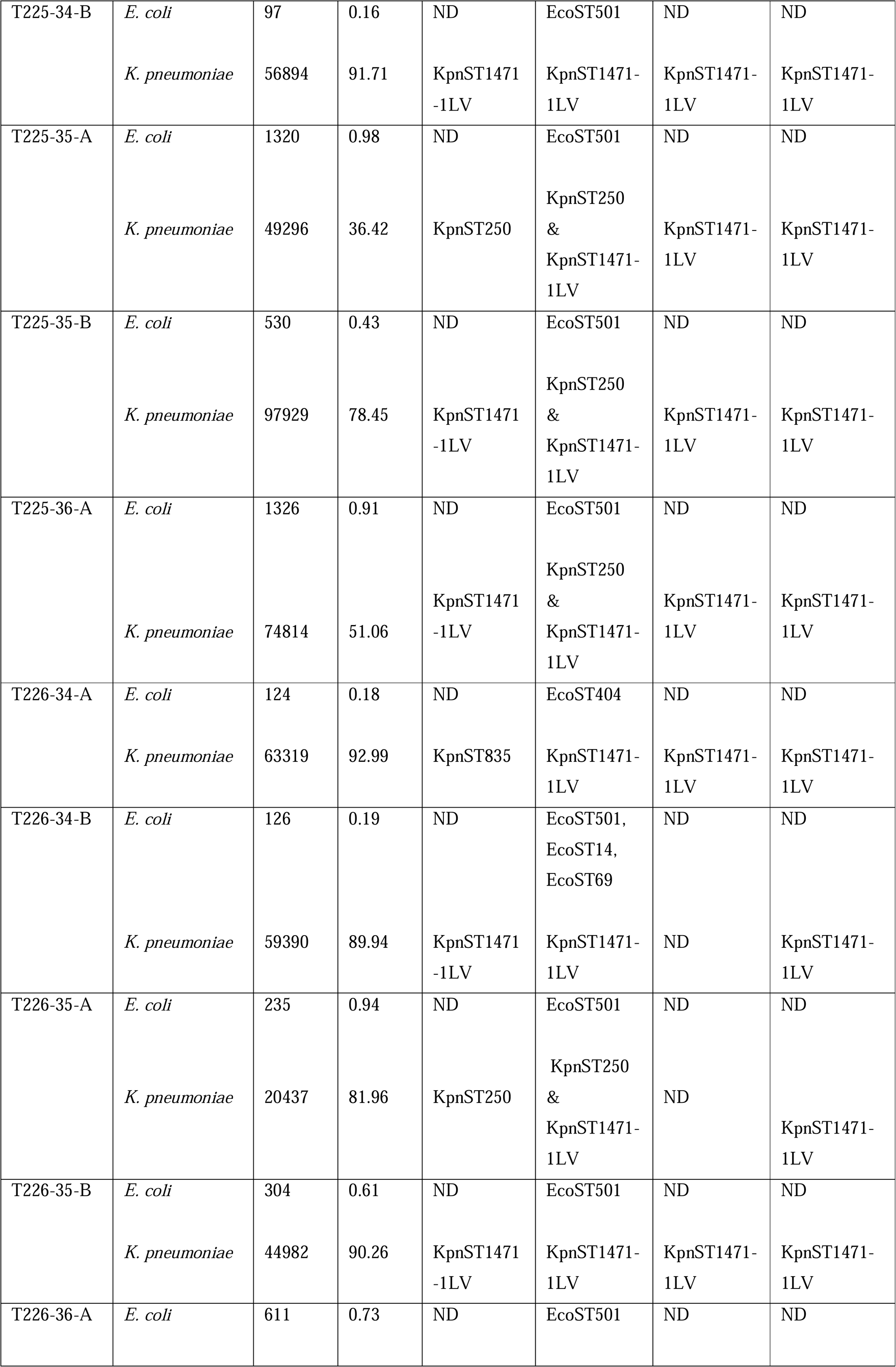
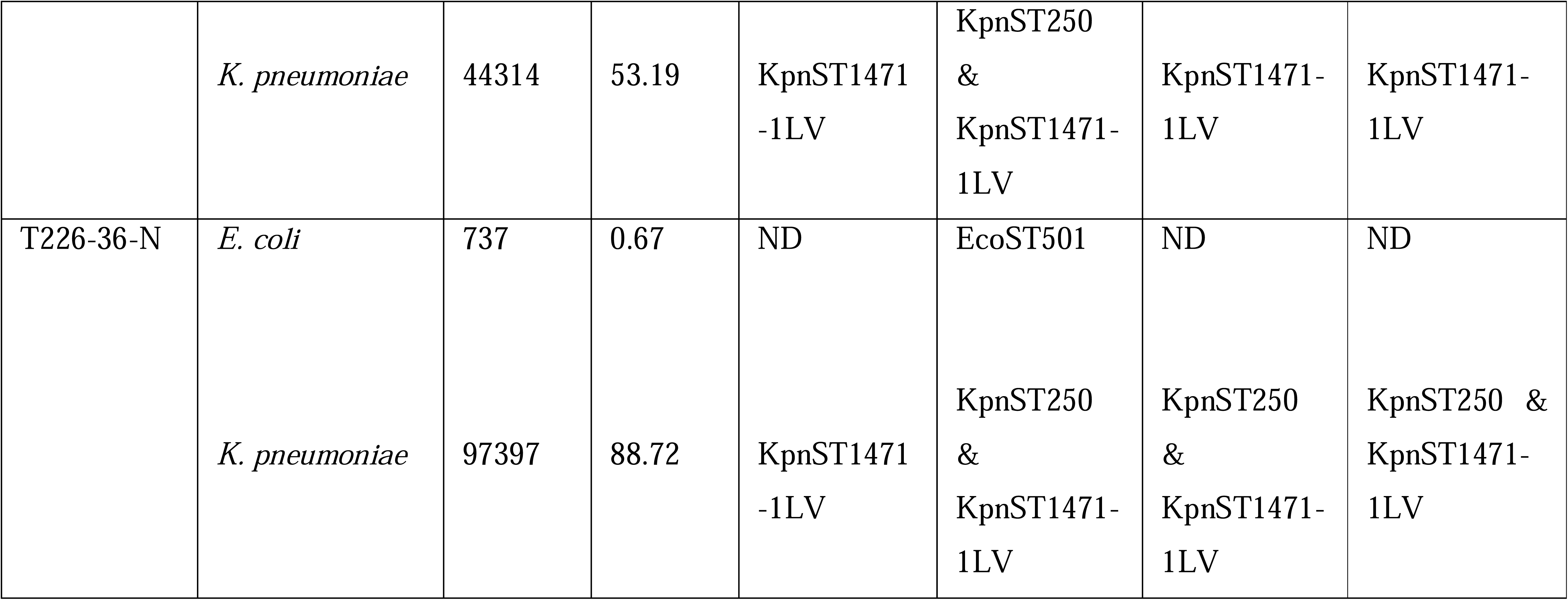
Comparison of strain resolution and taxonomic profile calls among TRACS, Strainy, and Strainberry. ND is an acronym for “Not detected”. In cases where at least one of the three tools identified dominant *E. coli* strains that were not detected through Illumina sequencing of pure cultures from clinical samples (T069-14-B, T098-12-B, T101-12-A, and T123-16-A), this discrepancy was likely due to technical oversight during the microbial culture process. Consequently, these samples were not preserved for re-culture and further analysis.

| Sample | Species<br>(Bracken) | Reads<br>(n) | Reads<br>(%) | Strain id<br>(Illumina) | Strain id<br>(TRACS) | Strain id<br>(Strainy) | Strain id<br>(Strainberry) |
| --- | --- | --- | --- | --- | --- | --- | --- |
| T069-12-A | *E. coli* | 69310 | 61.63 | EcoST8186<br>& EcoST95 | EcoST8186<br>& EcoST95 | EcoST8186 | EcoST8186,<br>EcoST95, &<br>EcoST95<br>(different<br>strain) |
|  | *K. pneumoniae* | 152 | 0.14 | ND | KpnST13 | ND | ND |
| T069-12-B | *E. coli* | 47345 | 44.17 | EcoST95 | EcoST95 | EcoST95 | EcoST95 |
|  | *K. pneumoniae* | 32 | 0.03 | ND | KpnST13 | ND | ND |
| T069-13-B | *E. coli* | 68562 | 95.56 | EcoST95 | EcoST95 | EcoST95 | EcoST95 |
|  | *K. pneumoniae* | 46 | 0.06 | ND | KpnST54 | ND | ND |
| T069-14-A | *E. coli* | 84562 | 92.92 | EcoST8186<br>& EcoST95 | EcoST8186<br>& EcoST95 | EcoST95 | EcoST95 |
|  | *K. pneumoniae* | 32 | 0.04 | ND | KpnST13 | ND | ND |
| T069-14-B | *E. coli* | 29001 | 68.37 | ND | EcoST95 | EcoST95 | EcoST95 |
|  | *K. pneumoniae* | 7 | 0.02 | ND | KpnST13 | ND | ND |
| T098-10-A | *E. coli* | 48052 | 98.52 | EcoST8186 | EcoST8186 | EcoST8186 | EcoST8186 |
|  | *K. pneumoniae* | 201 | 0.41 | ND | KpnST54 | ND | ND |
| T098-10-B | *E. coli* | 49409 | 99.32 | EcoST8186 | EcoST8186 | EcoST8186 | ND |
|  | *K. pneumoniae* | 5 | 0.01 | ND | KpnST54 | ND | ND |
| T098-11-A | *E. coli* | 60020 | 96.16 | EcoST8186 | EcoST8186 | EcoST8186 | EcoST8186 |
|  | *K. pneumoniae* | 147 | 0.24 | ND | KpnST54 | ND | ND |
| T098-11-B | *E. coli* | 56826 | 93.54 | EcoST8186 | EcoST8186 | EcoST8186 | ND |
|  | *K. pneumoniae* | 5 | 0.01 | ND | KpnST54 | ND | ND |
| T098-12-A | *E. coli* | 64074 | 97.23 | EcoST8186<br>& EcoST95 | EcoST8186 | EcoST8186 | EcoST8186 |
|  | *K. pneumoniae* | 6 | 0.01 | ND | KpnST54 | ND | ND |
| T098-12-B | *E. coli* | 30427 | 93.44 | ND | EcoST8186 | EcoST8186 | EcoST8186 |
|  | *K. pneumoniae* | 97 | 0.30 | ND | KpnST13 | ND | ND |
| T100-11-A | *E. coli* | 71605 | 97.65 | EcoST95 | EcoST95 | EcoST95 | EcoST95 |
|  | *K. pneumoniae* | 42 | 0.06 | ND | KpnST13 | ND | ND |
| T100-11-B | *E. coli* | 12045 | 94.35 | EcoST95 | EcoST95 | EcoST95 | EcoST95 &<br>EcoST95<br>(different<br>strain) |
|  | *K. pneumoniae* | 13 | 0.10 | ND | KpnST13 | ND | ND |
| T101-12-A | *E. coli* | 7206 | 97.68 | ND | EcoST95 | EcoST95 | ND |
|  | *K. pneumoniae* | 43 | 0.58 | KpnST104 | KpnST104 | ND | ND |
| T122-20-A | *E. coli* | 612 | 3.99 | ND | ND | ND | ND |
|  | *K. pneumoniae* | 6262 | 40.84 | KpnST1962 | KpnST1962 | ND | KpnST1962 |
| T123-16-A | *E. coli* | 13921 | 34.39 | ND | EcoST58 | ND | ND |
|  | *K. pneumoniae* | 24470 | 60.44 | KpnST5348 | KpnST5348 | KpnST5348 | KpnST5348 |
| T194-27-A | *E. coli* | 76392 | 76.88 | EcoST1722 | EcoST1722 | EcoST1722 | EcoST1722 |
|  | *K. pneumoniae* | 216 | 0.22 | ND | KpnST299 | ND | ND |
| T194-28-A | *E. coli* | 104350 | 92.58 | EcoST1722 | EcoST1722 | EcoST1722 | EcoST1722 |
|  | *K. pneumoniae* | 202 | 0.18 | ND | KpnST299 | ND | ND |
| T195-27-B | *E. coli* | 41614 | 63.91 | EcoST405 | EcoST405 | EcoST405 | EcoST405 |
|  | *K. pneumoniae* | 12020 | 18.46 | KpnST299 | KpnST299<br>(cluster 14) | KpnST299<br>(cluster 13) | KpnST299<br>(cluster 14) &<br>KpnST299<br>(cluster 14) |
| T225-34-A | *E. coli* | 144 | 0.20 | ND | EcoST69 &<br>EcoST1380 | ND | ND |
|  | *K. pneumoniae* | 66013 | 90.34 | KpnST1471<br>-1LV | KpnST1471-1LV | ND | KpnST 1471-1LV |
| T225-34-B | *E. coli* | 97 | 0.16 | ND | EcoST501 | ND | ND |
|  | *K. pneumoniae* | 56894 | 91.71 | KpnST1471-1LV | KpnST1471-1LV | KpnST1471-1LV | KpnST1471-1LV |
| T225-35-A | *E. coli* | 1320 | 0.98 | ND | EcoST501 | ND | ND |
|  | *K. pneumoniae* | 49296 | 36.42 | KpnST250 | KpnST250 & KpnST1471-1LV | KpnST1471-1LV | KpnST1471-1LV |
| T225-35-B | *E. coli* | 530 | 0.43 | ND | EcoST501 | ND | ND |
|  | *K. pneumoniae* | 97929 | 78.45 | KpnST1471-1LV | KpnST250 & KpnST1471-1LV | KpnST1471-1LV | KpnST1471-1LV |
| T225-36-A | *E. coli* | 1326 | 0.91 | ND | EcoST501 | ND | ND |
|  | *K. pneumoniae* | 74814 | 51.06 | KpnST1471-1LV | KpnST250 & KpnST1471-1LV | KpnST1471-1LV | KpnST1471-1LV |
| T226-34-A | *E. coli* | 124 | 0.18 | ND | EcoST404 | ND | ND |
|  | *K. pneumoniae* | 63319 | 92.99 | KpnST835 | KpnST1471-1LV | KpnST1471-1LV | KpnST1471-1LV |
| T226-34-B | *E. coli* | 126 | 0.19 | ND | EcoST501, EcoST14, EcoST69 | ND | ND |
|  | *K. pneumoniae* | 59390 | 89.94 | KpnST1471-1LV | KpnST1471-1LV | ND | KpnST1471-1LV |
| T226-35-A | *E. coli* | 235 | 0.94 | ND | EcoST501 | ND | ND |
|  | *K. pneumoniae* | 20437 | 81.96 | KpnST250 | KpnST250 & KpnST1471-1LV | ND | KpnST1471-1LV |
| T226-35-B | *E. coli* | 304 | 0.61 | ND | EcoST501 | ND | ND |
|  | *K. pneumoniae* | 44982 | 90.26 | KpnST1471-1LV | KpnST1471-1LV | KpnST1471-1LV | KpnST1471-1LV |
| T226-36-A | *E. coli* | 611 | 0.73 | ND | EcoST501 | ND | ND |
|  | *K. pneumoniae* | 44314 | 53.19 | KpnST1471<br>-1LV | KpnST250<br>&<br>KpnST1471-1LV | KpnST1471-1LV | KpnST1471-1LV |
| T226-36-N | *E. coli* | 737 | 0.67 | ND | EcoST501 | ND | ND |
|  | *K. pneumoniae* | 97397 | 88.72 | KpnST1471<br>-1LV | KpnST250<br>&<br>KpnST1471-1LV | KpnST250<br>&<br>KpnST1471-1LV | KpnST250 &<br>KpnST1471-1LV |

Furthermore, where TRACS detected the co-presence of two unique *E. coli* strains (EcST8186 and EcST95; 69310 – 84562 reads) and one *K. pneumoniae* strain (KpnST13; 32 – 152 reads) in two metagenomes, Strainy detected either an EcST8186 strain or EcST95 strain (Table 4; Figure 5B). On the other hand, Strainberry identified three *E. coli* strains (EcST8186, EcST95 (cluster_5), and EcST95 (cluster_11)) within 1 metagenome, and just one EcST95 strain in the other metagenome (Table 4; Figure 5B). Previous analysis using short-read sequence data from pure culture confirmed the presence of both EcST8186 and EcST95 strains in the same sample in agreement with TRACS’s analysis (Table 4).

Within a single metagenome where TRACS identified two other distinct *E. coli* strains (EcST69 and EcST1380; 144 reads in total) and one *K. pneumoniae* strain (KpnST1471-1LV; 66013 reads), Strainy failed to provide strain-level information. However, both Strainberry and short-read sequence data analyses confirmed the presence of the KpnST1471-1LV strain alone (Figure 6A, Table 4). Also, within one metagenome where TRACS identified the co-occurrence of three unique *E. coli* strains (EcST501, EcST14, and EcST69; 126 reads in total) and one *K. pneumoniae* strain (KpnST1471-1LV; 59390 reads), Strainy failed to detect any *E. coli* or *K. pneumoniae* strain (Figure 6A). On the other hand, Strainberry and short-read sequence data analyses confirmed the presence of the KpnST1471-1LV strain alone (Table 4).

**Figure 6:**
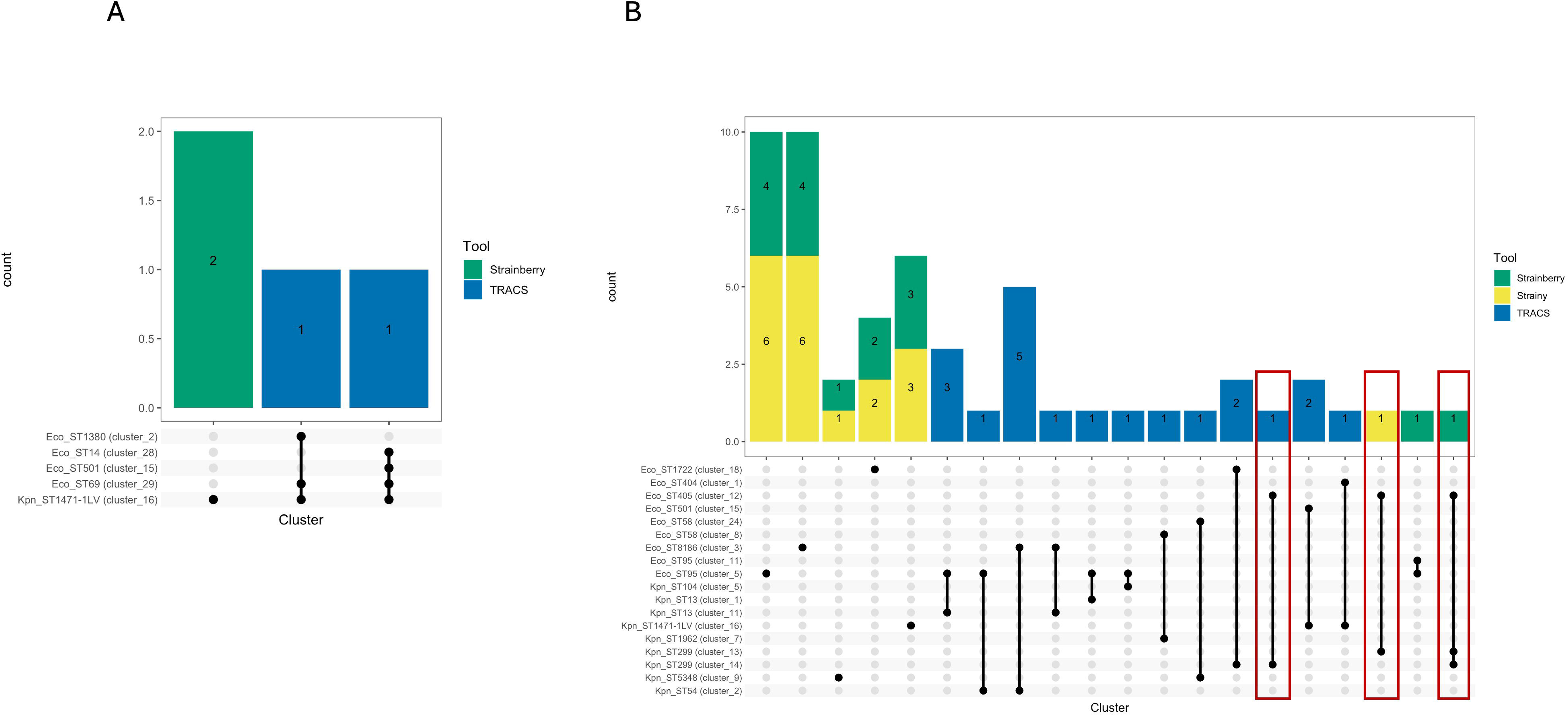
Upset plot illustrating combinations of strains identified/resolved by TRACS (blue colour), Strainy (yellow colour), and Strainberry (green colour) within real-world metagenomes, where TRACS identified more than one *E. coli* strain and one *K. pneumoniae* strain (A), or where TRACS identified one *E. coli* strain and one *K. pneumoniae* strain (B). The main bar chart displays the frequency of metagenomes per strain combination, arranged in descending order. Side bars indicate the number of metagenomes hosting each named strain. Dots and connecting lines at the base of the main bar chart represent the combination of *E. coli* and/or *K. pneumoniae* strains. Highlighted sections reveal differences in the resolution of the three tools when each of the tools resolved one *E. coli* strain and at least one *K. pneumoniae* strain from a metagenome.

In 20 of 21 other metagenomes where TRACS detected the concurrent presence of one *E. coli* strain and one *K. pneumoniae* strain, at least one strain of either organism was identified by either Strainy, Strainberry, or short-read data analysis (Figure 6B, Table 4). Interestingly, in one (containing Ec (41614 reads) and Kpn (12020 reads)) of the 21 metagenomes where TRACS identified the co-presence of one EcST405 strain and one KpnST299 strain (cluster 13), Strainy detected the co-presence of the identical EcST405 strain but a different KpnST299 strain (cluster 14) compared to TRACS’s identification, while Strainberry detected the presence of the identical EcoST405 strain alongside two KpnST299 strains, one from each of the clusters (clusters 13 and 14) identified by either TRACS or Strainy (red rectangular highlights to the right side of Figure 6B). Short-read sequence data analysis corroborated the presence of *E. coli* ST405 strain and *K. pneumoniae* ST299 strain (Table 4).

### TRACS demonstrates the highest haplotype completeness while Strainy demonstrates the lowest number of misassemblies in real-world metagenomic datasets

In our study, mirroring the methodology applied to mock metagenomic datasets, we utilized metaQUAST to assess the completeness and accuracy of strain assemblies from real-world metagenomic datasets. Notably, TRACS demonstrated the highest genomic fraction, achieving 99.9% for samples obtained from the anus and 99.8% for samples obtained from the nose of neonates. Strainy and Strainberry also exhibited notable genomic fractions, reaching up to 96.2% and 97.8% for samples from the anus, and 81.2% and 85.8% for samples from the nose, respectively.

Strainy demonstrated the lowest number of misassemblies, with a maximum of 82 and 41 misassemblies observed in samples from the anus and nose, respectively. Conversely, TRACS and Strainberry displayed higher misassembly counts, with up to 110 and 98 misassemblies in anal samples, and 68 and 92 misassemblies in nasal samples, respectively.

### TRACS demonstrates the highest process execution speed and computational resource efficiency except single-core CPU usage

TRACS demonstrated efficiency by completing the alignment of real-world metagenomes against a custom reference database and resolving strains in just 22 minutes. In contrast, Strainy and Strainberry took significantly longer, requiring 78 minutes and 291 minutes, respectively, to resolve strains from metagenomes (Figure 7A). TRACS also showcased better resource efficiency, utilizing the lowest amount of physical memory compared to Strainy and Strainberry (Figure 7B). Notably, the higher execution time and higher memory usage of Strainberry and Strainy can be attributed to their dependence on metagenomic assembly to generate a strain-oblivious metagenome before strain resolution. Nevertheless, Strainy used the lowest percentage of single-core CPU usage at 113%. Strainberry exhibited the highest percentage of single-core CPU usage at 331%, while TRACS utilized 176% of single-core CPU (Figure 7C).

**Figure 7:**
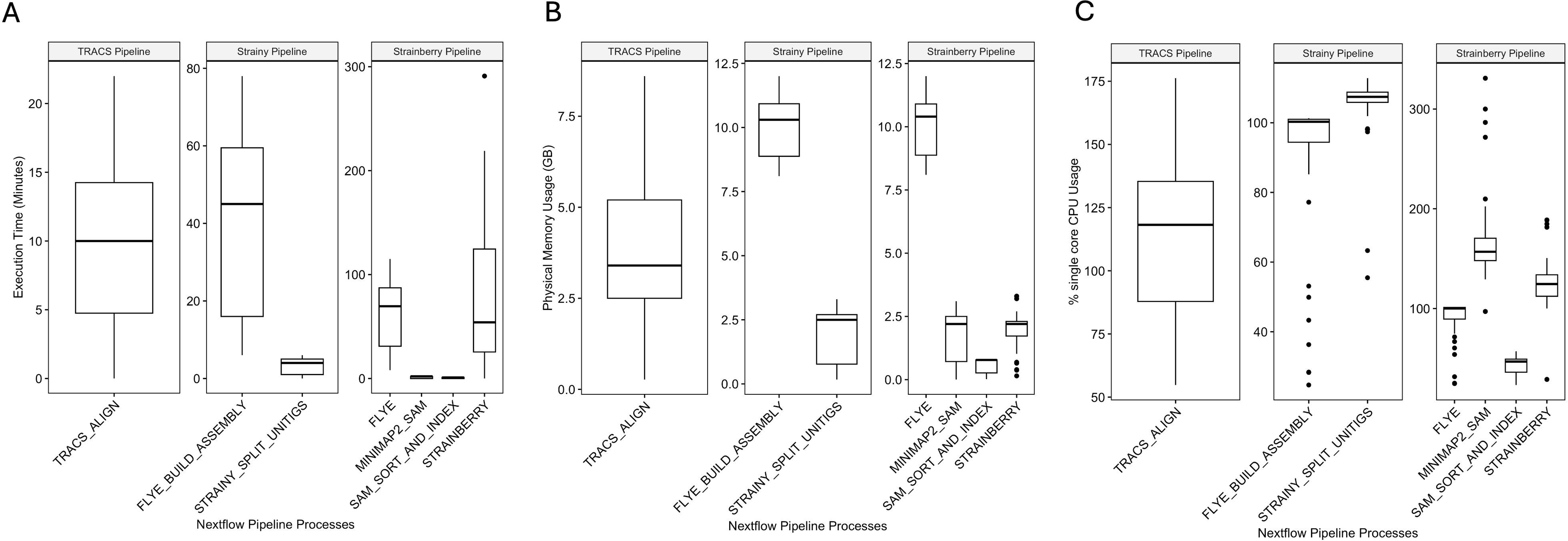
Assessment of the computational performance of the three strain resolution tools, including execution time (A), physical memory usage (B), and percentage single-core CPU usage (C).

## Discussion

In this study, we evaluated the performance of three strain resolution tools, TRACS, Strainy, and Strainberry, using mock community datasets and real-world metagenomic datasets derived from the Oxford Nanopore Technology platform. Furthermore, we assessed execution time, memory usage, and CPU utilization of each tool to elucidate their efficiency in resolving strains.

The evaluation of strain resolution tools on mock communities revealed comparable performance among TRACS, Strainberry, and Strainy. The performance remained consistent across different flow cell chemistries and dominant species within the mock community. Notably, Strainberry tended to overestimate the number of strains present in the mock communities while Strainy tended to underestimate the number of strains present in the mock microbial community. In general, all strain resolution tools were unable to identify all strains when the mock community contained strains from the same sequence type. This suggests that while the tools are effective in detecting strain diversity, they may have limitations in accurately quantifying strain numbers, particularly in communities with high genetic similarity among strains.

The observed differences in haplotype completeness and accuracy among TRACS, Strainberry, and Strainy are consistent across mock community datasets and real-world datasets. While TRACS demonstrated higher haplotype completeness, Strainy exhibited superior accuracy, emphasizing the trade-offs between genome fraction and misassembly rates. These findings align with existing literature [13] and underscore the importance of considering both metrics as well as other metrics in evaluating the performance of strain resolution tools.

Analysis of real-world metagenomic datasets highlighted the ability of TRACS, Strainy, and Strainberry to resolve strains within clinical samples, to detect strains that had previously gone undetected in short-read sequencing of pure cultures, and to confirm microbiologically negative samples. For instance, all three strain resolution tools correctly reported the absence of *E. coli* and *K. pneumoniae* in microbiologically negative samples. However, Bracken assigned a very low number of reads to these species despite the samples being classified as microbiologically negative. This phenomenon is consistent with previous benchmarking studies, which indicate that highly sensitive metagenomic classifiers, such as Bracken and Kraken, may produce low-abundance false positives [32]. Specifically, certain methods have been shown to call false positives at abundances below 0.5%, especially when analyzing complex communities or low-diversity samples [32]. In this study, the low-abundance signals from Bracken likely reflect background noise rather than true biological presence, in line with the known microbiological status of the samples as negative. This observation highlights the importance of selecting appropriate abundance cutoffs when interpreting metagenomic data to minimize false-positive classifications.

Additionally, there were instances where all three tools successfully detected dominant *E. coli* and/or *K. pneumoniae* strains (both in terms of read count and relative abundance within the metagenome) that had not been identified through traditional microbiological methods. In some cases, subsequent re-culturing from stored samples confirmed the presence of these strains. However, in other cases, culturing (particularly of *E. coli* strains) was not performed due to technical oversights during sample handling, and as a result, the original swab samples were not stored and short-read sequencing of pure cultures could not be conducted. This emphasizes the added value of metagenomic analysis for strain detection—especially in scenarios where traditional cultivation methods/processes might be incomplete or overlooked [33, 34].

While each tool demonstrated strengths in detecting strains, discrepancies were observed in certain instances which may be attributed to differences in their underlying approaches. While Strainberry employs an iterative pairwise phasing algorithm for variant calling and haplotype phasing from error-prone long-read sequence data aligned against metaFlye assemblies, Strainy utilizes a different phasing approach for variant calling, haplotype phasing and strain identification from the alignment of long-read sequence data against metaFlye assembly graphs. Both tools rely on MetaFlye because it yields the best assemblies in terms of total metagenome size and metagenome recovery from nanopore-based metagenomic sequencing, compared to other known long-read assemblers such as Canu, Raven, Readbean, or Shasta [35]. In contrast, TRACS employs a kmer-based approach to infer pairwise transmission from single genomes or raw metagenomic sequence data. These variations in methodology, particularly in the alignment step versus kmer-based approaches, may account for differences in results and highlight the importance of understanding the underlying algorithms when interpreting strain resolution outcomes. In a clinical setting, however, it is critical to select the tool that demonstrates the best overall performance in terms of accuracy and reliability, as this will likely capture the majority of the relevant strains, even if some may be missed. This approach ensures efficient and consistent strain-level resolution that is actionable for clinical decision-making.

TRACS was most efficient in process execution speed and computational resource utilization compared to Strainy and Strainberry. TRACS completed the alignment and strain resolution of real-world metagenomes in significantly less time and with lower physical memory usage. However, Strainy demonstrated the lowest single-core CPU usage, while Strainberry exhibited the highest, suggesting greater computational demands and highlighting differences in resource utilization strategies among the tools. Notably, the higher execution time of Strainberry and Strainy can be attributed to their dependence on metagenomic assembly to generate a strain-oblivious metagenome before strain resolution, which contributes to increased processing time and memory usage. These distinctions in computational efficiency have broader implications beyond laboratory time and cost; they directly impact the environmental footprint associated with high-performance computing in genomics. By reducing CPU time, memory usage, and execution time, TRACS minimizes the energy consumption associated with large-scale metagenomic analyses, thereby contributing to a lower carbon footprint. Given the growing adoption of metagenomics in clinical and research applications, optimizing computational workflows to reduce resource demands will play an essential role in supporting both sustainable research practices and environmental stewardship [36].

Our analysis of real-world metagenomic datasets revealed several challenges encountered by strain resolution tools, particularly when applied to complex microbial communities and utilizing real databases. While our efforts aimed to maintain consistency in the composition of the reference database used for evaluating the performance of strain resolution tools across mock community datasets and real-world datasets, we acknowledge that hurdles encountered with authentic datasets may stem from limitations inherent in the tools and reference database employed. Additionally, discrepancies in strain identification and assembly completeness underscore the need for continuous refinement and optimization of strain resolution algorithms, as well as the development of comprehensive reference databases tailored to specific clinical and research contexts. Addressing these challenges will be crucial for advancing our understanding of microbial community dynamics and improving the reliability of strain-level analyses in clinical and research settings. Finally, determining the minimum threshold of reads required to reliably resolve strains poses a significant challenge. In the real-world metagenomes analysed in this study, instances occurred where only a small number of reads were attributed to a specific strain. Remarkably, in many of these cases, at least one tool, often TRACS, effectively identified the strain, aligning with the findings from short-read sequence data analysis in at least one scenario. However, in other cases, TRACS may have been calling false positives when read numbers were low, particularly as it consistently identified certain strains, such as KpnST13 and KpnST54 (for *Klebsiella pneumoniae*; both STs do not belong to the same clonal complex) and EcoST501 (for *Escherichia coli*), in low-read scenarios where both other tools and short-read sequence data analysis from pure cultures detected no such strains. In a few instances, TRACS even detected multiple strains of *E. coli* or *K. pneumoniae* in these low-read cases, such as identifying three strains of *E. coli* from only 126 reads. Nevertheless, in microbiologically negative samples where a substantial number of reads (more than 200) were allocated to a specific strain, all three tools correctly detected the absence of an *E. coli* or a *K. pneumoniae* strain. Given that read count and depth are critical factors influencing haplotype completeness (which is particularly challenging to achieve for low-abundance organisms), establishing the minimum read depth necessary for accurate strain resolution is crucial for practical applications, especially in optimizing multiplexing and shortening long-read sequencing time. Understanding these thresholds can inform experimental design choices and guide sequencing strategies, ultimately enhancing the efficacy, environmental sustainability, and cost-efficiency of genomic analyses.

In conclusion, our study provides valuable insights into the performance and limitations of strain resolution tools in microbial community analysis. By comprehensively assessing these tools on mock and real-world datasets, we contribute to the ongoing efforts to improve the accuracy and reliability of strain-level resolution in clinical and research settings. Moving forward, continued optimization and benchmarking of strain resolution algorithms are essential to enhance our understanding of microbial community dynamics and facilitate more informed clinical decision-making.

## Funding information

The work was funded by the Federal Ministry of Education and Research (BMBF) under 01KI2018 to SR.

## Author contributions

Conceptualization: S.R. Data curation: A.O.A., S.A.M., I.J.A., and L.D.d.S. Data Analysis and Investigation: A.O.A. Project administration and supervision: S.R. Resources: S.R. Visualization: A.O.A. Writing – original draft: A.O.A. Writing – review & editing: All authors.

## Conflicts of interest

The authors have no conflicts of interest to declare.

## Abbreviations

ST: sequence type
NICU: neonatal intensive care unit
ONT: Oxford Nanopore Technology
GTDB: Genome Taxonomy Database
MAG: metagenome-assembled genome
SNP: single nucleotide polymorphism

## Data Summary

Mock metagenomic sequence datasets and real-world metagenomic sequence datasets have been submitted to the ENA under bioprojects ID PRJEB82665 and PRFEB82667 (Table S1).

## Impact Statement

Our research addresses a critical challenge in microbial genomics and public health by evaluating the performance of three bioinformatics tools—TRACS, Strainy, and Strainberry—in resolving strains within clinical microbial communities. By employing advanced long-read sequence-based methods, we demonstrate the potential of these tools to overcome limitations in strain resolution, thus expanding their applicability beyond specialised genomic laboratories equipped with state-of-the-art sequencing technologies and expert bioinformaticians. These laboratories typically perform complex genomic analyses that require significant expertise and resources, which are often not available in routine clinical or public health settings. By systematically evaluating TRACS, Strainy, and Strainberry, this work identifies key differences in accuracy, computational efficiency, and genome completeness, providing critical insights into the capabilities of each tool. TRACS exhibited a better computational efficiency and strain completeness, offers a promising solution for resource-limited settings or high-throughput applications. This study contributes valuable insights into improving microbial surveillance and outbreak response strategies, ultimately advancing our ability to combat infectious diseases and safeguard public health.

---

**Supplementary Figure 1:** Maximum-likelihood phylogenetic tree of core genomes depicting microbial strains within the mock microbial community. Known *K. pneumoniae* strain identities are juxtaposed with strains detected by the three strain resolution tools. Annotations on the right side of the tree denote strains’ sequence type as well as expected and predicted mock composition, with red-shaded circles indicating presence and empty circles indicating absence. Black dots within red-shaded circles signify strains belonging to the same cluster (SNP < 20) as the expected strain identity. Section A and section B refer to mock community comprising up to two strains belonging to different sequence types and same sequence types, respectively. Section C and section D refer to mock community comprising up to three strains belonging to a different sequence type and same sequence types, respectively.

**Supplementary Figure 2:** Evaluation of strain-resolved assemblies generated from mock metagenomic sequencing data analysis by the three strain resolution tools using metaQUAST. The figure depicts the assembled strain genome fraction and the number of misassemblies for each *E. coli* strain in the mock metagenome datasets. Strains are arranged in decreasing genome fraction value among all tools.

**Supplementary Figure 3:** Evaluation of strain-resolved assemblies generated from mock metagenomic sequencing data analysis by the three strain resolution tools using metaQUAST. The figure depicts the assembled strain genome fraction and the number of misassemblies for each *K. pneumoniae* strain in the mock metagenome datasets. Strains are arranged in decreasing genome fraction value among all tools.

**Supplementary Table 1:** ENA id and Bioproject id of mock community datasets and real-world metagenomic datasets.

## Notes

### Competing Interest Statement

The authors have declared no competing interest.

