## Supplementary figures and images for "Assessing the performance of current strain resolution tools on long-read metagenomes"

### Supplemental Figure 1

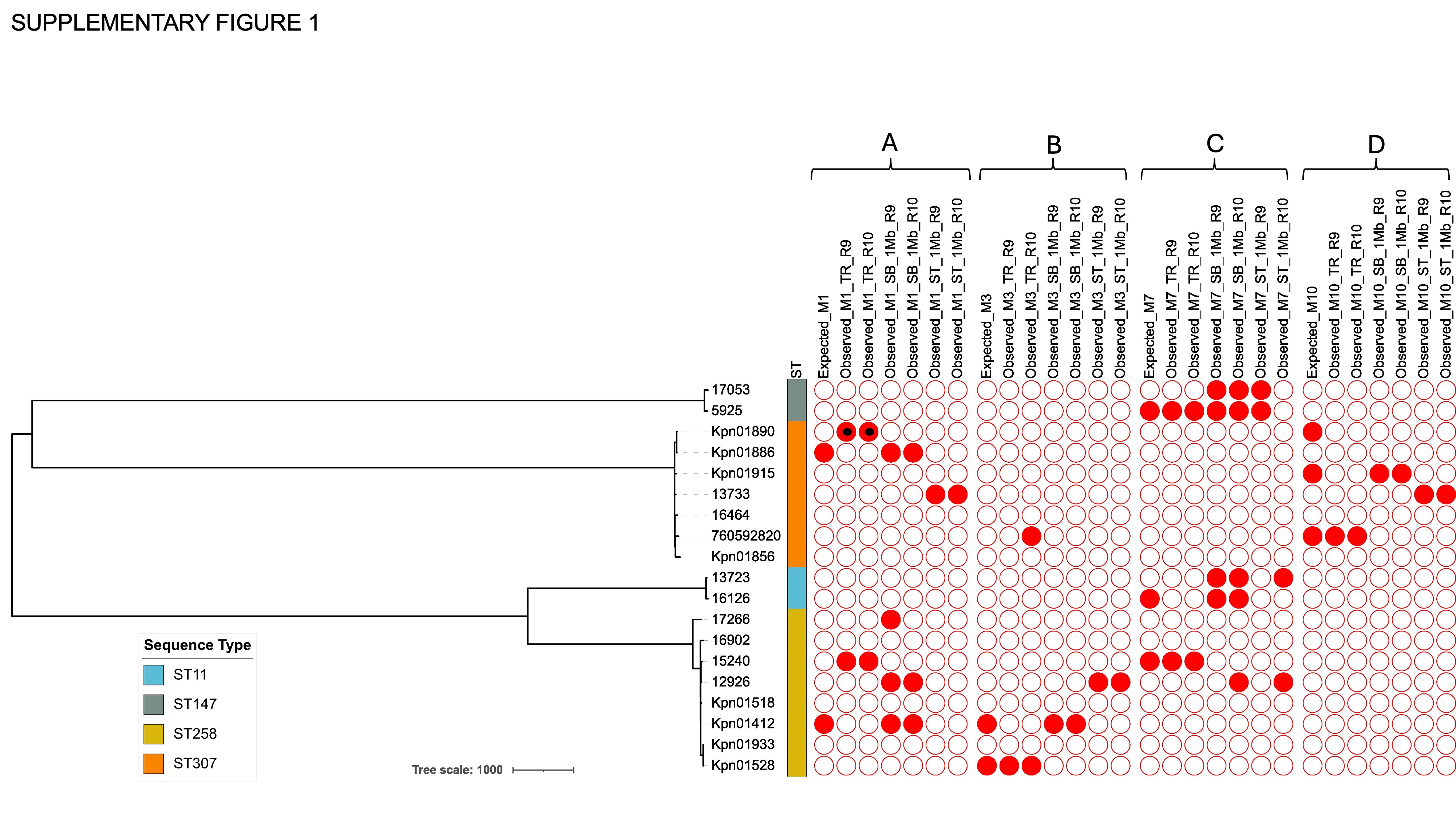

### Supplemental Figure 2

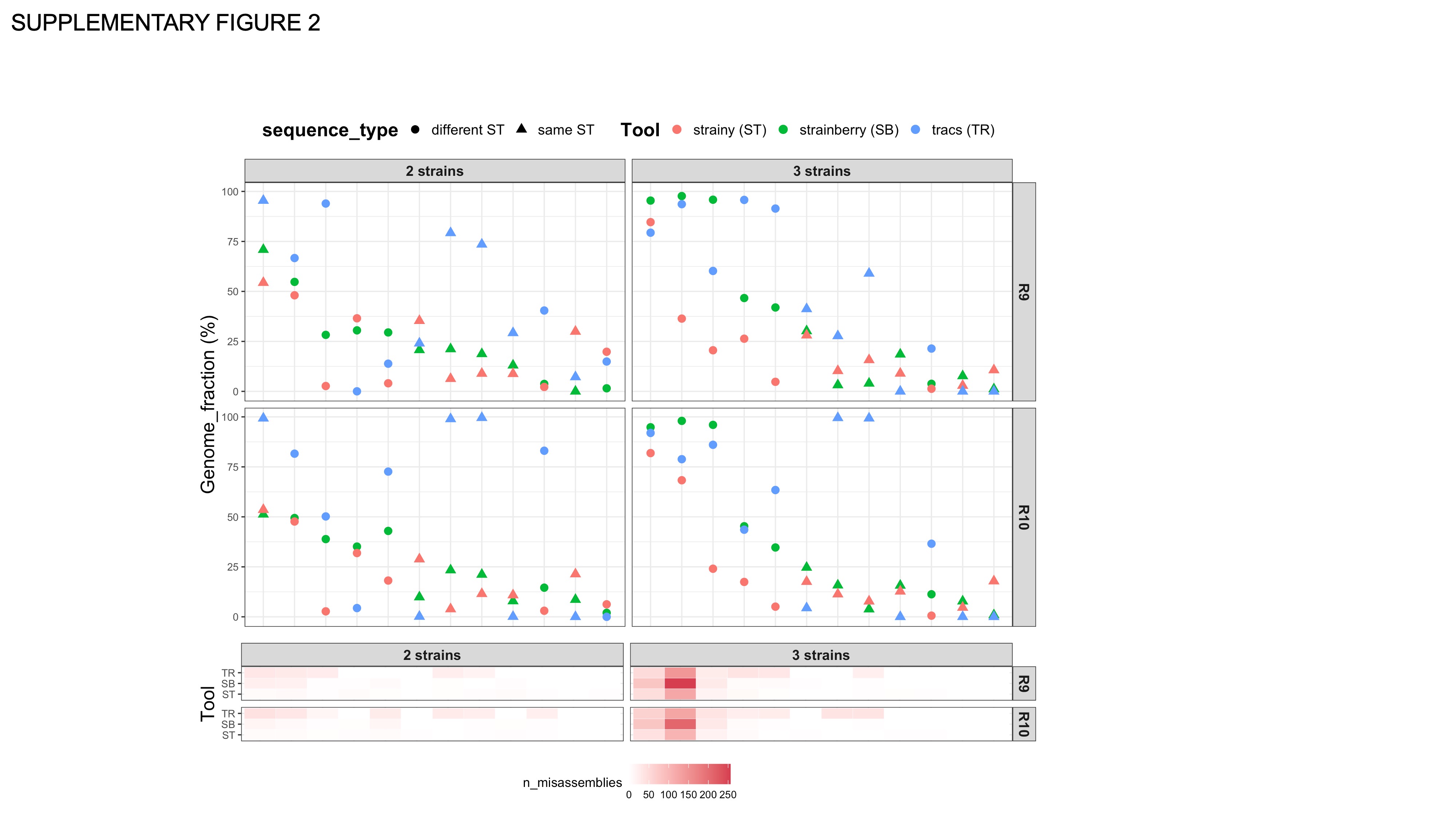

### Supplemental Figure 3

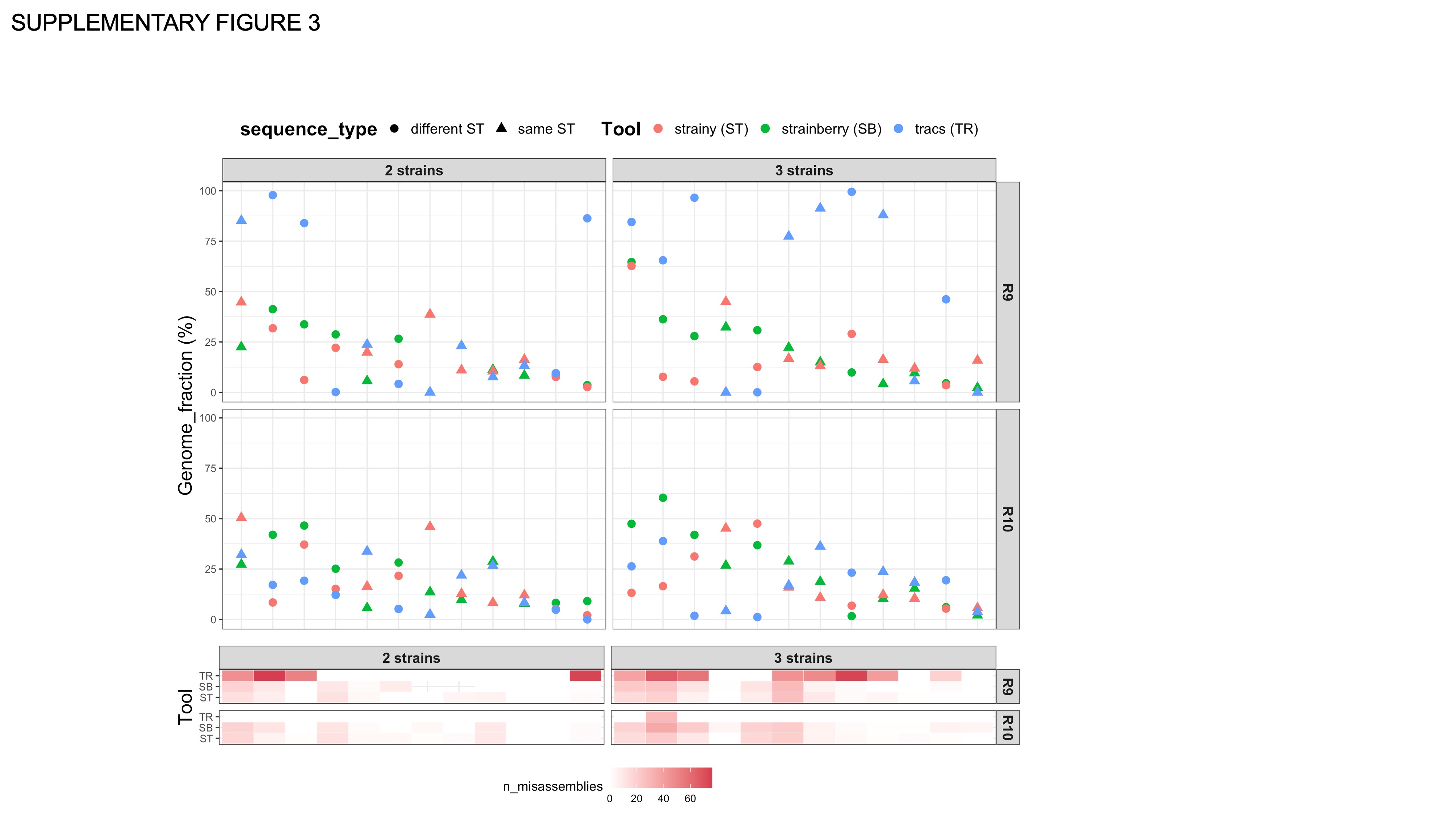
